# Explaining decision biases through context-dependent repetition

**DOI:** 10.1101/2024.10.09.617399

**Authors:** Ben J. Wagner, H. Benedikt Wolf, Stefan J. Kiebel

## Abstract

Humans are prone to decision biases, which make behavior seemingly irrational. An important cause for decision biases is that the context in which decisions are made can later influence which choices humans prefer in new situations. Current computational models often require extensive environmental knowledge to explain these biases. Here, we tested the hypothesis that decision biases are mainly driven by a tendency to repeat context-specific actions. We implemented a series of nine value-based decision experiments on n = 351 male and female participants and reanalyzed six previously published datasets (n=350 participants). We found that higher within-context repetition of an option was associated with biased choices including higher subjective valuation and lower uncertainty for repeated actions. Next, we used a computational model based on two basic principles, learning by reward and decision repetition and tested it on all datasets. Our results show that the combination of these two basic principles is sufficient to explain biased choices. We demonstrate via hierarchical Bayesian model comparison that our model outperforms all alternative models. These results provide novel insights into human choice behavior and offer a solution to difficulties in current explanations and models.

**Significance Statement:** Human decision-making often deviates from rational choice theory. Understanding how decision biases emerge is essential for interpreting real-world human behavior. Our study shows how repeating decisions shapes decision making in novel situations. We adapt a simple computational model that can explain decision biases based on reward learning and decision repetition. By testing the model across 15 datasets, we demonstrate that it outperforms previous models, also offering new insights into how preferences affect subjective valuation and uncertainty. In sum, these findings provide a deeper understanding of how context-dependent preferences for specific decisions emerge and persist.

## Introduction

Unlike an unconstrained rational agent, human information processing is limited by memory capacity and processing bandwidth (Bhui et al. 2021). Contextualized processing might be one adaptive solution to such limitations as it constrains the stream of information, i.e. what memories or action plans are accessed (Heald et al. 2023). Evidence for the role of context in human cognition includes classical and instrumental conditioning (Courville et al. 2006), the interplay of context and memory in extinction learning (Gershman et al. 2010; Bouton et al. 2021), the role of context in associative-, structure- or representation learning (Gershman 2017; Rosas et al. 2013; Niv 2019), the context-specific interplay of habitual and goal-directed behavior (Wood and Rünger 2016; Schwöbel et al. 2021; Schwöbel et al. 2024; Froelich et al. 2024) or contextual inference in motor learning (Heald et al. 2021; Cuevas Rivera and Kiebel 2023). Experimentally, strong effects of context have also been shown in value-based decision-making tasks where participants first learn to make choices between rewarding options and subsequently show counter-intuitive decision biases (Klein et al. 2017; Bavard et al. 2018; Bavard et al. 2021; Bavard and Palminteri 2023; Molinaro and Collins 2023b). Specifically, in the first phase of such experiment’s participants learn the value of different options in exclusive choice contexts. For example, participants must make repeated choices between two options A and B (in context one) and options C and D (in context two), but never between options across contexts (e.g., A and C). After this learning phase, participants enter a so-called transfer phase. In this phase participants make choices between two options across contexts, i.e. choices between new combinations of options such as A and C, or between B and D. It is in this transfer phase that participants often make choices that appear surprisingly counter-intuitive or even irrational, especially when viewed through the lens of expected utility theory, which assumes that humans behave as if they maximize subjective utility. For instance, humans often display a strong and systematic choice preference when none should exist, such as when both A and C should have the same expected value (Klein et al. 2017; Bavard and Palminteri 2023; Glimcher 2022; Molinaro and Collins 2023b). In some cases, participants even prefer losses over gains (Bavard et al. 2018).

In recent years various mechanisms and computational models have been proposed to explain these counter-intuitive preferences. Some of these models were inspired by the discovery of normalization of neuronal signals, such as contrast gain control in the retina (Shapley and Victor 1978) or auditory cortex (Rabinowitz et al. 2011). In consequence it has been suggested that normalization might be a key principle that also applies to how the brain stores and retrieves value representations (Louie and Glimcher 2012; Louie et al. 2015). Further research investigated which specific form of normalization explains human choices best (Bavard et al. 2021; Bavard and Palminteri 2023) or whether humans balance between internal and external reward signals, given an internal goal to obtain the maximum outcome (Molinaro and Collins 2023a, 2023b). One potential issue with these approaches is that they in principle require specific statistical knowledge about the environment. For example, for range normalization (Bavard and Palminteri 2023) or the weighting of intrinsic and extrinsic reward signals (Molinaro and Collins 2023b) participants are assumed to update value estimates with respect to the outcomes of all alternatives. However, such knowledge may or more often may not be available in real-world decision making.

We propose a computationally simpler account of these biased preferences, based on standard reinforcement learning (RL) and a tendency to repeat previous context-specific choices. In this framework, consistent with Thorndike’s *Law of Exercise* (Thorndike 1911), we suggest that merely making a choice can bias the likelihood of selecting that option in the same context again. Several experimental and modeling studies present evidence that preference formation may be shaped by both reward-based and repetition-driven mechanisms, i.e. a repetition bias (Lau and Glimcher 2005; Seymour et al. 2012; Akaishi et al. 2013; Katahira et al. 2017; Miller et al. 2019; Gershman 2020; Schwöbel et al. 2021; Palminteri 2023; Legler et al. 2024; Schwöbel et al. 2024; Lai and Gershman 2024; Froelich et al. 2024), and that making and repeating a choice might be related to its valuation (Brehm 1956; Sharot et al. 2009; Enisman et al. 2021). Theoretically, this repetition bias may arise from cognitive resource constraints, as perseveration-like behavior can serve as an adaptive strategy to reduce cognitive effort in complex decision environments (Gershman 2020; Bhui et al. 2021; Lai and Gershman 2024).

To comprehensively test such a context-specific repetition-bias mechanism, we collected nine new datasets (n = 351 participants) and conducted a reanalysis of six datasets from published studies (n = 350 participants). We demonstrate that the proposed mechanism accounts for biased preference across all tasks. These include tasks involving probabilistic and continuous rewards, losses and gains, tasks with as few as two or as many as 66 across context comparisons, and tasks where feedback is provided either for all alternatives or only for the selected option. Using formal model comparison based on hierarchical Bayesian modeling, we compare this model to all previously proposed alternatives: relative value learning (Klein et al. 2017), divisive normalization (Louie et al. 2011; Louie and Glimcher 2012; Webb et al. 2021), variants of range normalization (Bavard & Palminteri, 2023b) and a model that weights external and internal reward signals (Molinaro and Collins 2023b). In summary, our findings show that the proposed repetition-bias (REP) model is the best explanation across almost all analyzed datasets (n = 701 participants). Moreover, it is the only model that implements a mechanism for the correlation between choice frequency during learning and choice preference during transfer, observed across all datasets. Additionally, our results show that relative choice frequency of an option is linked to both the perceived value of this option and the certainty about that value. Participants showed greater certainty about options they had selected more frequently, even when full feedback was available and tended to overvalue these frequently chosen options compared to those selected less often. We further discuss the implications of our findings for a re-conceptualization of decision-biases.

## Results

### Standard analyses

We first perform a descriptive assessment of the choice data during learning and test whether participants show any systematic preference during the beginning of the transfer phase. We then present three key results from our standard analyses across all 15 datasets: (i) there is a significant correlation between participants’ relative choice frequencies in the learning phase and their choice preferences in the transfer phase; (ii) the relative choice frequency during learning is linked to post-task valuation of options, specifically, options chosen more frequently are overvalued compared to those chosen less frequently; and (iii) the relative choice frequency also relates to participants’ certainty about an option’s value, with participants being more certain about the value of options they have chosen more often during learning.

### Assessment of preferences

In those tasks, where both available contexts differed by contrast (p1 and p2 or g1 and g2) participants on average chose the best option, i.e. the option with the highest expected value, C_HC_(0.7) or D_HC_(0.7)) in the high contrast (HC; see Figure 1A) context more often than the best option (A_LC_(0.7)) in the low contrast context (LC; see Figure **2A-C & F-G**). Similarly, and as expected, during transfer, most participants did prefer the best HC option (C_HC_ in probabilistic tasks or D_HC_ in Gaussian tasks) over its equally valued counterpart (A_LC_). We here report exact binomial tests to examine whether the proportion of choosing C_HC_ or D_HC_ in the first transfer trial (TT) is not equal to 0.5. (see Figure **2H** and **I** for task p1 (full feedback; p < 0.0001; CI[0.63-0.82]) and p2 (partial feedback; p = 0.08; CI[0.48-0.77]) and Figure **2M** and **N** and for task g1 (full feedback; p < 0.001; CI[0.59-0.85]) and g2 (partial feedback; p < 0.001; CI[0.60-0.86])). Strikingly, in task p3 (Figure **2J**), this relationship reverses when the LC context was encountered more often during learning (50 times vs. 30 times) and thus the best LC option was by design chosen more often than the best HC option (p = 0.004; CI[0.16-0.43]). This shows that the HC effect is marginalized by a higher number of repetitions. In task 4.1, where participants chose options, during learning, in approximately the same relative choice frequency (green and orange lines in Figure **2D**), clear choice preference during transfer is absent, even if expected values favor a high gain (HG) option over a low gain (LG) option (Figure **2K;** p = 1; CI[0.31-0.69]). Likewise, in task 4.2 (Figure **2E** and **L**), choice preference during transfer is clearly visible (p < 0.001; CI[0.64-0.94]; Figure **2L**), when participants had a clear favorite option during learning, as measured by within context relative choice frequency (Figure **2E**). In terms of probabilistic tasks, effects were larger when full feedback was provided and diminished in the partial feedback task. This can be seen in Figure2**A** (full feedback) and **B** (partial feedback) for the average within context relative choice frequencies during learning, i.e. red (C_HC_) and blue (A_LC_) lines. This difference is numerically more pronounced when full feedback is provided (Figure **2A**). Likewise, the preference for this HC option (red; C_HC_) over the LC option (blue; A_HC_) during transfer is more pronounced in the full-(p1: p < 0.0001; CI[0.63-0.82]) compared to partial feedback task (p2:; p = 0.08; CI[0.48-0.77]; see Figure **2H & I**).

**Figure 1:**
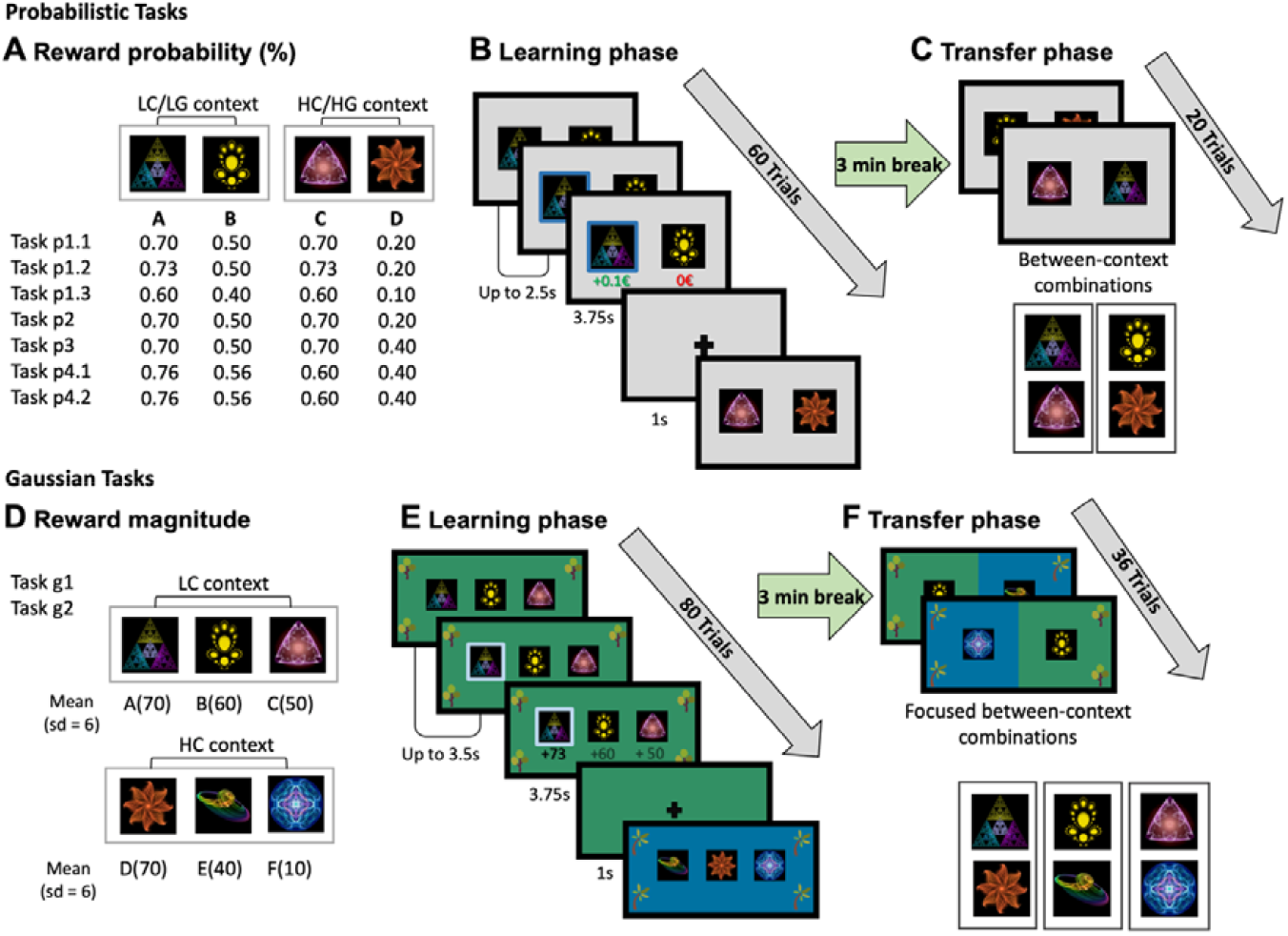
Overview of value-based decision-making tasks (acquired data). Top row: Probabilistic Tasks. **A:** Reward probabilities for all four options (A-D), encountered in both the low contrast (LC) and high contrast (HC) choice contexts in tasks p1.1 to p3, and in the low gain (LG) and high gain (HG) choice contexts in tasks p4.1 and p4.2. **B:** Learning phase for probabilistic tasks. Participants are presented with pairs of options (abstract fractal stimuli) within both LC/LG and HC/HG contexts and must select one option per trial. Across multiple trials, they learn to choose the better option through trial-and-error. In tasks p1.1–p1.3 and p3–p4.2, full feedback was provided for both the chosen and unchosen options. In task p2, only partial feedback was provided for the chosen option. Trials are presented in a pseudorandomized order, ensuring that the actual reward probabilities are maintained across every 10–12 trials (for exact sequences, see Supplemental Table 5). For analysis purposes, tasks p1.1 to p1.3 were pooled into a single dataset (task p1) due to their shared features and the absence of significant differences in results when analyzed individually. **C:** In the transfer phase, options from both LC/LG and HC/HG choice contexts are tested against each other to evaluate whether participants show systematic biases, such as a preference for the C_HC_(0.70) option over the A_LC_(0.70) option. **D:** Reward magnitudes for Gaussian tasks. In these tasks, trial-wise reward magnitudes for each option are sampled from a Gaussian distribution with a standard deviation of six. Each choice context consists of three options. In the LC context, reward distributions for different options do overlap, making it more challenging to identify the best option. **E:** Learning phase for Gaussian tasks, equivalent to **B**. Participants make repeated choices between options within the LC and HC choice contexts. To help distinguish between contexts, visual cues are provided: distinct background colors (blue or green) and different tree types (evergreen or broadleaf trees) displayed in the four corners of the screen. **F:** In the transfer phase, equivalent to **C**, participants’ preferences across contexts, options from both the LC and HC choice contexts are tested against each other. This mixed-context presentation is indicated by combined visual cues, including a mix of background colors and tree types. In all tasks, participants’ cumulative monetary reward was calculated by summing their trial outcomes: €0.10 for each correct choice in probabilistic tasks, or an amount proportional to the points earned in Gaussian tasks (0 to 100 points multiplied by €0.01). Participants received their cumulative reward as a bonus payment upon completing each task.

**Figure 2:**
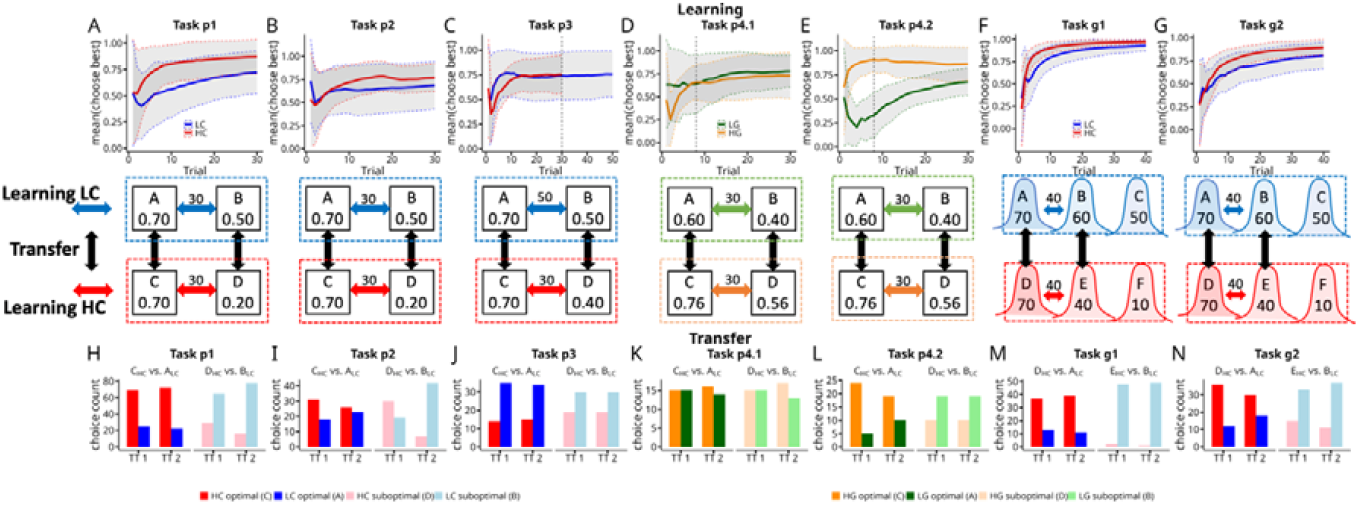
Relative choice frequency (within context) during learning and choice preference during transfer. **Top row (A-G):** Relative choice frequency (within context) during learning across all probabilistic and Gaussian tasks. **A:** Task p1 (full feedback), relative choice frequency for the best option in both the low contrast (LC; blue) and high contrast (HC; red) contexts. **B:** Task p2 (partial feedback), relative choice frequency for the best option in the LC (blue) and HC (red) contexts. **C:** Task p3 (full feedback), relative choice frequency for the best option in LC (blue) and HC (red) contexts. **D:** Task p4.1 (full feedback), relative choice frequency for the best option in low gain (LG; green) and high gain (HG; orange) contexts. **E:** Task p4.2 (full feedback), relative choice frequency for the best option in LG (green) and HG (orange) contexts. **F:** Task g1 (full feedback), relative choice frequency for the best option in LC (blue) and HC (red) contexts. **G:** Task g2 (partial feedback), relative choice frequency for the best option in LC (blue) and HC (red) contexts. **Middle row (A-G):** Task parameters and context repetition counts. **A:** Task p1 (collapsed p.1 to p.3), reward probabilities and number of repetitions of LC and HC contexts (30 trials for each context during learning). In the LC context, option A_LC_ had a 0.70 reward probability, and option B_LC_ 0.50. In the HC context, option C_HC_ had a 0.70 and D_HC_ a 0.20 reward probability. **B-E:** Tasks p2, p3, p4.1, p4.2, reward probabilities and number of repetitions of each context, as in A. **F**: Task g1, reward magnitudes for Gaussian tasks, represented by points. In the LC context, options A, B, and C offered average rewards of 70, 60, and 50 points, respectively. In the HC context, options D, E, and F offered 70, 40, and 10 points. **G:** Task g2, analogous to panel F but with partial feedback. **Bottom row (H-N):** Choice counts for available options within the first and second transfer trial (TT), illustrating participants’ choice preferences when options from different contexts are presented together. **H**: Task p1, absolute preference in TT 1 and TT 2 for the best option in the HC context (red, option C_HC_) and the optimal option in the LC context (blue, option A_LC_) as well as for the options with lower reward probabilities (B_LC_ (pink) and D_HC_ (light blue)). **I:** Choice counts in TT1 and TT2 in task p2 (partial feedback). **J:** Choice counts in TT1 and TT2 in task p3 (full feedback). Here, participants more often chose the best LC option (A_LC_) indicating an effect of repetition during learning. **K:** Choice counts in TT1 and TT2 in task p4.1 (full feedback). **L:** Choice counts in TT1 and TT2 in task p4.2 (full feedback). **M:** Choice counts in TT1 and TT2 in task g1 (full feedback). **N:** Choice counts in TT1 and TT2 in task g2 (partial feedback).

### Frequency correlation between learning and transfer

We next analyzed whether relative choice frequency (across contexts) during learning and relative choice preference during transfer (within context) are significantly correlated. To do this, we computed, for the learning phase, relative choice frequencies for all pairwise combinations of options encountered in the transfer phase. For example, in task p1 (Figure **2A & H**), we calculated how often participants chose, during learning, option C_HC_ relative to how often they chose option A_LC_ (relative frequency learning; see methods). Note that by design options C_HC_ and A_LC_ were never paired in the same trial during learning, but one can compute their relative choice frequencies, i.e., how often each option was selected in relation to the other. Analogously, for the transfer phase, we computed how often participants preferred C_HC_ over A_LC_ during the first two transfer trials when directly compared against each other (relative frequency transfer; see methods). We found that relative choice frequency during learning and relative choice preference during transfer was highly and significantly correlated across all datasets (pooled data across all probabilistic tasks incl. Klein et al. 2017; r = 0.80, p = 0.0002), Gaussian tasks (ρ = 0.98, p < 0.0001), both datasets from Bavard et al. 2018 (Exp1: r = 0.72, p < 0.0001; Exp2: ρ = 0.83, p < 0.0001) and two datasets from Bavard et al. (2023; Exp3a: ρ = 0.92, p < 0.0001; Exp3b: ρ = 0.86; p < 0.0001). However, computing correlations across all pairwise combinations can be problematic, as these correlations may be influenced by expected reward. For example, in tasks p1 and p2, participants might choose option B_LC_(0.5) more frequently than option D_HC_(0.2) during learning, solely because B_LC_(0.5) has a higher expected value. Similarly, if successfully learned, participants may prefer option B_LC_(0.5) over D_HC_(0.2) during transfer because of the difference in expected value. To address this issue, we aimed to control for such effects by analyzing three different subsets of the data: First, we selected only those option pairs from all 15 datasets where both options had equal expected values (for exact selection criteria, see methods). We still found a strong and significant correlation between choice frequency during learning and choice preference during transfer (ρ = 0.66, p < 0.001; Figure **3A**). The presence of this correlation indicates that participants’ choice preference for an option during transfer is related to the relative choice frequency for that option during learning (relative to the compared transfer option with equal expected value). Crucially, expected value cannot be an explanation for this correlation because all compared options had the same expected value and were not directly compared during learning. Second, one might argue that even though objective expected values of the compared options were equal, preferences could still be influenced by relative values, i.e. implying that humans learn values on a relative scale, i.e. as assumed by state-dependent normalization (Klein et al. 2017; Bavard et al. 2018). To address this possibility, we computed context-specific relative values for each option by normalization and selected only those option pairs from all 15 datasets where both options had the same normalized values (for exact selection criteria, see methods). We found that the relative choice frequency during learning and the relative choice frequency during transfer remained strongly correlated (ρ = 0.67, p = 0.002; see Figure 3B). This further indicates that frequency plays a role in preference formation. If frequency had no effect on preference, or if participants relied exclusively on state-dependent normalization, we would expect no correlation in this case. Third, to provide evidence that loss preference during transfer, as reported by Bavard et al. (2018), might as well be related to choice frequency during learning, we selected all option pairs from their study where participants had to choose between gains and losses of varying magnitudes during the transfer phase. Analysis of these selected data revealed a strong correlation between choice frequency during learning and choice preference during transfer (r = 0.88, p < 0.0001; see Figure **3C**). Importantly, when an option associated with a loss (small loss, relative loss, or big loss) had a higher choice frequency relative to an option associated with a gain during transfer, we found that such a loss option always had a higher relative choice frequency during learning (all red datapoints above 0.5 (y-axis; relative frequency transfer) fall within the upper-right green rectangle).

**Figure 3:**
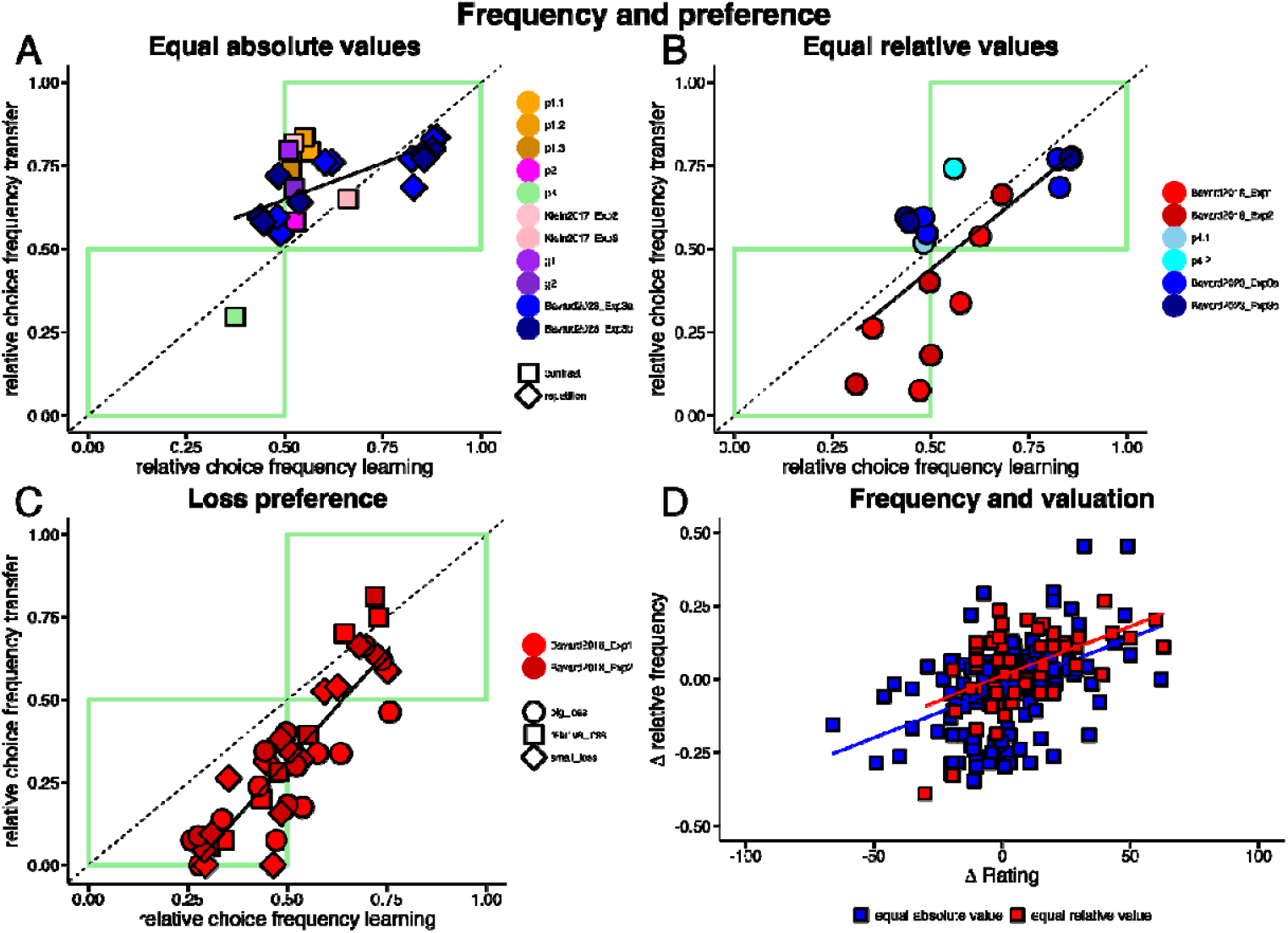
**A-C** Linear relationships between relative choice frequency during the learning phase (x-axis; relative frequency learning) and preference during the transfer phase (y-axis; relative frequency transfer) for selected pairwise combinations of options participants encountered in the transfer phase. Data points (colored dots; colors indicate different datasets) represent relative choice frequencies, see methods. Green rectangles: Highlights data points where an above (upper right) or below (lower left) 50% relative choice frequency, during learning, is associated with an above or below 50% relative choice frequency during transfer. **A:** Relative choice frequences of option pairs with equal expected values. Nearly all pairwise combinations fall within the upper right green rectangle, indicating that options preferred during the transfer phase (>50%) were chosen more frequently than their counterpart (>50%) during the learning phase. Correlation analysis showed a strong correlation of relative choice frequency during learning and transfer (ρ = 0.66, p < 0.001). **B:** Relative choice frequencies of option pairs with equal relative values. Results indicate that even when relative values are equal, relative choice frequency during learning is related to relative choice frequency (preference) in transfer (ρ = 0.67, p = 0.002). **C:** Loss preference: Option pairs where losses are compared against gains, from Bavard et al. (2018). This panel illustrates that preferences for losses or smaller gains (relative losses) during transfer follow the same positive relative frequency learning – relative frequency transfer relationship (r = 0.88, p < 0.0001). Data points are differentiated by shape according to three categories to illustrate specific influences on decision-making: Small loss category: Comparisons between an option associated with a small loss (−0.1€) and options associated with a gain (+0.1€, +1€). X-values are relative frequencies for the small loss when normalized with respect to the compared gain option. Relative loss category: Comparisons between an option associated with a small gain (0.1€) and options associated with a big gain (+1€). X-values represent relative choice frequencies for the small gain option when normalized with respect to the compared large gain option. Big loss category: Comparisons between an option associated with a big loss (−1€) and a small or big gain (0.1€, 1€). X-values are relative frequencies for the big loss when normalized with respect to the compared small or big gain option (see Bavard et al. 2018 for further details). All loss-associated options preferred during the transfer phase (as indicated by their placement within the upper right green rectangle) were chosen more frequently (above 50%) during learning than their higher-valued counterparts. Options outside the green rectangles do not necessarily violate the linear relationship between relative choice frequency during learning and transfer; instead, they likely reflect the dominance of expected value effects over frequency. **D**, Relationship between differences in normalized choice frequencies (Δ relative frequency; see methods) and post-task option valuation (Δ rating; difference in rating between both options). Data points are color-coded based on whether the choices involved only equally valued options (blue, C_HC_ vs. A_LC_ in probabilistic tasks or D_HC_ vs. A_LC_ in Gaussian tasks) or options with equal relative value (red, C_HG_ vs. A_LG_ from tasks p4.1 an p4.2). This plot shows how differences in normalized choice frequency of two options correlate with differences in participants’ post-task valuation of the two options. The positive association of differences in normalized frequency and option valuation indicates that valuation is positively related with choice frequency.

### Frequency and valuation

If higher relative choice frequency of an option causes preference for that option during the transfer phase, one would also expect this to be reflected in participants’ valuation of options. Specifically, we computed relative choice frequencies at the participant level and assessed whether differences in relative choice frequency between two options were associated with differences in their post-task valuation (see methods for details). To control for the obvious influence of reward, we focused our analysis as above to pairwise comparisons of options with equal expected values. In tasks with such pairs (p2, p3, g1, g2), we found that differences in relative choice frequency were significantly correlated with differences in post-task valuation. This correlation was robust whether relative choice frequency was computed from the combined learning and transfer phase data (r = 0.43, p < 0.00001; blue squares in Figure **3D** or from the learning phase data alone (r = 0.42, p < 0.00001). We next focused on options with equal relative value (tasks p4.1 and p4.2; C_HG_(0.76; +0.1 relative value) and A_LG_(0.6; + 0.1 relative value)) and likewise computed the correlation between differences in relative choice frequency and differences in post-task valuation. Again, we found significant positive correlations when using combined learning and transfer phase data (r = 0.51, p < 0.0001; red squares in Figure **3D**) and learning phase data only (r = 0.47, p < 0.0002). These findings indicate that participants’ relative choice frequency is linked to their post-task valuation of options. When directly compared, options chosen more frequently were overvalued compared to those chosen less frequently.

### Frequency and uncertainty

Similarly, we analyzed whether relative choice frequency is related to uncertainty about the value of options. On the group level the spread of the belief distribution (standard deviation) about option value is significantly and negatively correlated with relative choice frequency in probabilistic tasks (p2, p3, p4.1 and p4.2; ρ = −0.647 =, p = 0.0082; Figure **4G**) and Gaussian tasks (g1 and g2; ρ = −0.90 p <0.0001; Figure **4F**). In addition, on the individual participant level higher relative choice frequency was related to lower distance from the true expected value (probabilistic tasks) or from the mean of the Gaussian reward magnitudes (Gaussian tasks) when correlated across all participants and tasks (r= −0.1 p = 0.006).

**Figure 4:**
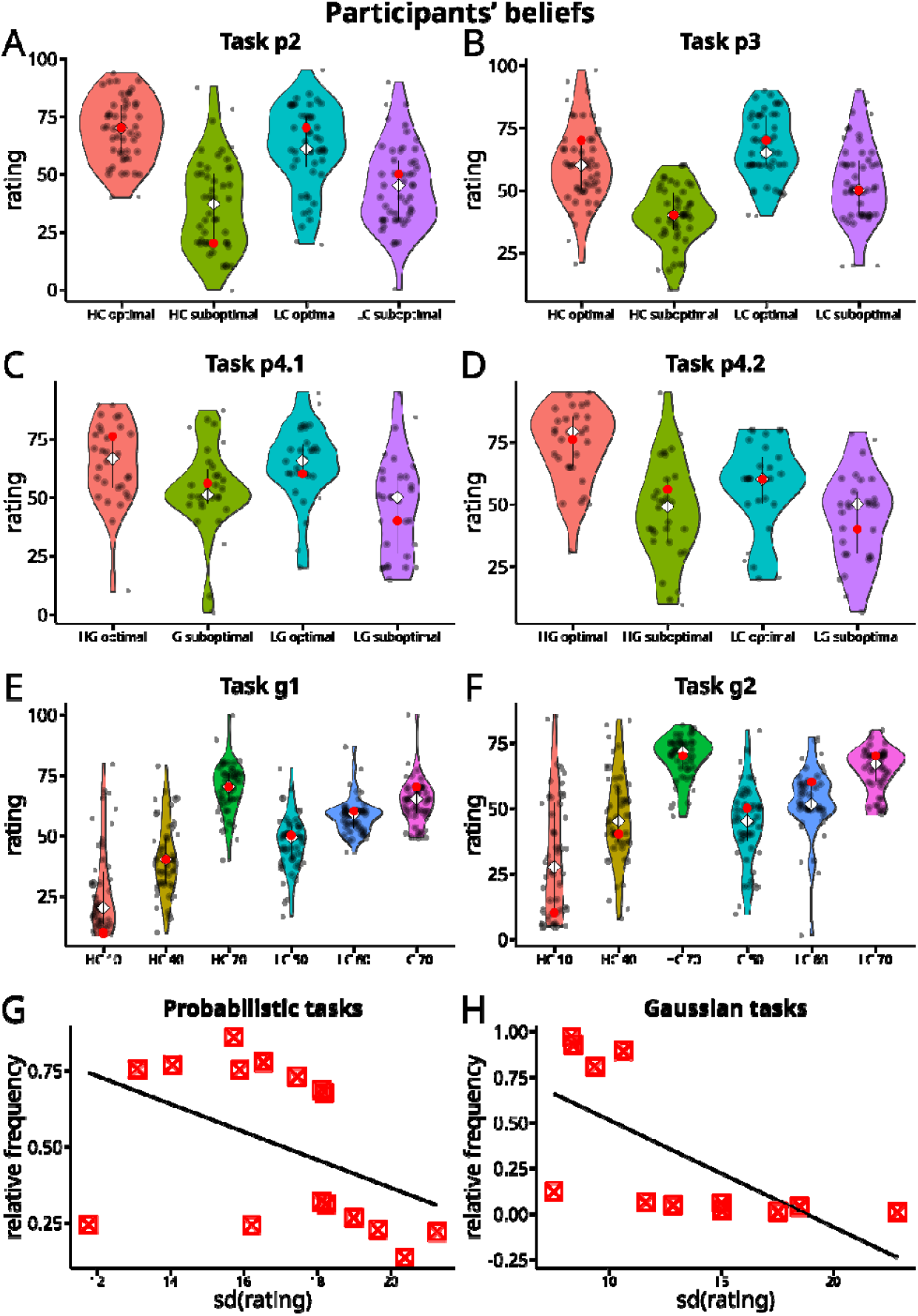
Post-task valuation ratings and relationship between uncertainty and choice frequency (data was not collected for task p1). **A–F** Violin plots for post-task valuation ratings (subjective beliefs) with true reward probabilities or reward magnitudes of each option (red dots), mean of participants’ beliefs about these task parameters (white diamonds), and individual participants’ ratings of each option (black dots). **A:** Valuation ratings for task p2 (n = 49). **B:** Valuation ratings for task p3 (n = 49), **C:** Valuation ratings for task p4.1 (n = 29). **D:** Valuation ratings for task p4.2 (n = 30), **E:** Valuation ratings for task g1 (n = 50). **F:** Valuation ratings for task g2 (n = 49). **G:** Relationship between participants’ (group level) uncertainty about an option and normalized choice frequency in the learning phase, for all probabilistic tasks except p1. The standard deviation (SD) of participants’ individual valuation ratings as a measure of belief uncertainty is plotted on the x-axis. Normalized choice frequency of each option during the learning phase is plotted on the y-axis. We found a significant negative correlation (ρ = –0.647, p = 0.0082). **H:** Equivalent to G but for Gaussian tasks (g1 and g2) with a significant negative correlation (ρ = –0.90, p < 0.0001).

### Computational modelling

To better understand the mechanisms driving the correlation between choice frequency during learning and choice preference during transfer, we analyzed choice data from all tasks using a computational model and compared its performance against all prominent alternative models (Louie and Glimcher 2012; Klein et al. 2017; Bavard and Palminteri 2023; Molinaro and Collins 2023b). The proposed model integrates two key components: (i) standard reinforcement learning (RL) and (ii) reward-independent learning of a context-specific repetition bias (Akaishi et al. 2013; Katahira et al. 2017; Schwöbel et al. 2021; Palminteri 2023; Froelich et al. 2024). For the first component, we assumed participants learn Q-values for each option in an unbiased manner as would be assumed in standard RL (Sutton & Barto, 2018). We here tested two different model variants. For the first we used different decaying learning rates for each context (Sutton and Barto 2018), and for the second, following previous studies (Palminteri et al. 2015; Bavard et al. 2018; Bavard and Palminteri 2023) different learning rates for chosen and unchosen options (see methods for details). For the second component, we let choices increase or decrease the repetition bias of an option:

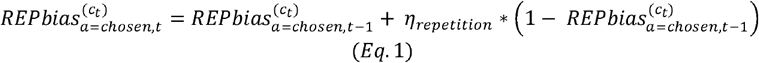

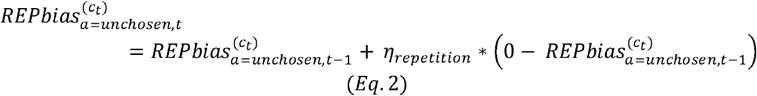

where the repetition-bias (REPbias) for each option of the current context is updated using a separate learning rate parameter *η*_*repetition*_. The REPbias for an option increases on a trial-wise basis when an option is chosen (Eq. 1) and decreases when the option is not chosen (Eq. 2). Using hierarchical Bayesian modeling and quantitative model comparison, we tested this model (called in the following the repetition (REP) model) against all recently proposed computational mechanisms to explain choice preferences in the transfer phase. From the literature, we identified six different models. These are relative value learning (REL model; Klein et al. 2017), divisive normalization (DIV model; Louie and Glimcher 2012), variations of range normalization (RANGE & RANGE models; Bavard and Palminteri 2023), the intrinsic enhancement model (IER model) proposed by Molinaro and Collins (2023b) and a fully unbiased base model where Q-values are learned on an absolute scale (ABS model) (see Supplemental Table 1 for an overview). For datasets based on probabilistic tasks with constant reward magnitude (tasks p1, p2, p3, p4.1, p4.2 and Klein et al. 2017) we compared the REP, ABS, REL and IER models (subset 1; see methods for details on model selection). For Gaussian tasks and probabilistic tasks with changing reward magnitude (g1, g2, Bavard et al. 2018 and Bavard and Palminteri 2023) we compared the REP, ABS, DIV, RANGE, RANGE*ω* and IER models (subset 2; see methods). In the following we focus on model comparisons using context specific decaying learning rates. Results for models with constant learning rates for chosen and unchosen options were mostly similar and are reported in the supplement.

### Probabilistic tasks (new datasets)

First we analyzed task p1 (full feedback) and task p2 (partial feedback) which show the well-established preference for an option of equal value (see Figure **2A & B** and **H & I**) when it has been learned in a high contrast (HC) context, compared to an alternative learned in a low contrast (LC) context (e.g., see Klein et al. (2017), Bavard et al. 2018). As expected, the ABS model cannot qualitatively predict these findings. Theoretically, the REL model should predict this outcome because the best option in the HC context (0.5 value difference; see task parametrizations in Figures 1 and 2) has a higher relative value (+0.25) compared to the best option in the LC context (0.2 value difference; +0.1 relative value). The IER model should explain this preference due to less counterfactual feedback, which should increase the intrinsic enhanced signal for the best HC option (Molinaro & Collins, 2023b). The REP model should also account for this preference as the best HC option was chosen more frequently, thereby increasing the REPbias for this option. For task p1 (pooled data for p1.1-p1.3; for separate model fits see Supplemental Table 3), we find that quantitative model comparison using the deviance information criterion (DIC; Spiegelhalter et al. 2014) favored the REP model (DIC: 4213.49, lower values indicate better fit) compared to the REL (DIC: 4339.53), ABS (DIC: 4562.59) and IER (DIC: 4562.51) models, see Figure **5A**. For task p2 (partial feedback) the results are qualitatively similar, i.e. participants showed a consistent preference for the best option learned in the HC context. Again, the REP model (DIC: 2696.46) performed better than the ABS (DIC: 2835.23), REL (DIC: 2902.83) or IER (DIC: 2848.28) models (see Figure **5A**).

Task p3 was explicitly designed to test whether a preference during transfer for one of the two highly rewarded options (A_LC_(0.7) or C_HC_(0.7)) was driven either by reward centering (as in the REL model), by an intrinsic signal (as predicted by the IER model), or by choice frequency, i.e. repetition (as considered by the REP model). To test this, we increased the number of LC context repetitions relative to the number of HC context repetitions (see Figure **2C**; 50 times vs 30 times). In theory, both the REL and IER models will predict a preference for the best HC option (C_HC_) because there was less counterfactual feedback or it had a higher relative value. In contrast, the REP model predicts a preference for the best LC option because this option was chosen more often during the learning phase due to the increased number of LC trials. The behavior shown during the transfer phase supported the REP model’s prediction as participants showed a strong preference for the best LC option (see Figure **2H**). Congruent with observed behavior during transfer, model comparison shows that the REP model had the lowest DIC (DIC: 3503.70), outperforming the IER (DIC: 3653.67), REL (DIC: 3584.49), and ABS (DIC: 3650.67) models (see Figure **5A** and Supplemental Table 3). In tasks p4.1 and p4.2, both contexts in each task featured options with an equal expected value difference of 0.2 but they differed in their absolute expected values resulting in a low gain (LG) and high gain (HG) context (LG context: A_LG_ (0.60) vs. B_LG_ (0.40) and HG context: C_HG_ (0.76) vs. D_HG_ (0.56); for task parametrizations and experimental results see Figure **2D & E** and **K & L**). Here, we focused on a potential choice preference between option A_LG_ (0.60) and option C_HG_ (0.76). In theory, the REL model would predict indifference for either A_LG_ or C_HG_ because both options are associated with equal relative value. The other models do not make specific predictions, in fact their predictions vary depending on exact parametrizations. We found that the dataset for task p4.1 was again best explained by the REP model (DIC: 1591.80); REL (DIC: 1724.28); ABS (DIC: 1794.51); IER (DIC: 1796.2). Task data for p4.2 was best explained by the REL model (DIC: 1364.51), with the REP model being a close second (DIC: 1386.69). The other two models had a higher DIC: IER (DIC: 1438.14) and ABS (DIC: 1439.19); see Figure **5A**) Note that standard analyses speak against the REL model, because participants showed a clear transfer preference and no evidence of indifference (see Figure **2E**).

### Gaussian tasks (new data)

Next, we present the results of our Gaussian tasks g1 and g2, where we tested the REP model against several normalization models: divisive normalization (DIV), range normalization (RANGE), weighted range normalization (RANGEω), and the IER model. According to the theory underlying range normalization, participants should be indifferent between the A_LC_(70) and D_HC_(70) options because both options would be normalized equally when considering the context-specific ranges (LC context: A(70)-B(60)-C(50) and HC context: D(70)-E(40)-F(10)). Contrary to this prediction, participants showed a clear preference for the D_HC_(70) option over the A_LC_(70) option in both the full and partial feedback tasks (g1 and g2; see Figure **2M & N**). For task g1, the IER model provided the best fit (DIC = 2074.69), followed by the REP model (DIC = 2183.07) and the RANGEω model (DIC = 2264.18). The other models performed worse, with DIC values of 2595.10 (ABS), 3042.93 (RANGE) and 3602.51 (DIV). For task g2, the RANGEω model provided the best fit (DIC = 4243.95), followed by the REP model (DIC = 4314.60) and the IER model (DIC = 4356.02). The other models performed worse with DIC values of 4451.66 (ABS), 4480.37 (RANGE) and 5403.30 (DIV).

### Previously published datasets

To further assess the generalizability of the REP model, we conducted a comparative model analysis using data from previously published studies that introduced alternative models. First, Klein et al. (2017), whose datasets are in principle equally parametrized as our tasks p1 and p2, found evidence in favor of the REL model (when compared to the ABS model). Our reanalysis, as in the analyses of the new datasets p1 and p2, confirmed the original finding that the REL model (DIC k_exp2: 1710.31; k_exp3: 1494.54) explained decision biases in the transfer phase better than the unbiased model (DIC(ABS) exp2: 1767.75; exp3: 1542.66). However, when including other models into the comparison, the REP model had the lowest DIC (DIC(REP) exp2: 1527.09; exp3: 1426.89), outperforming both the REL and IER models (DIC(IER) exp2: 1767.71; exp3: 1555.91; see Figure **5B** and Supplementary Table 3). The Bavard et al. (2018) dataset allowed us to test the REP model in a task that included both gain and loss contexts, different magnitudes, and mixed partial and full reward feedback. The REP model performed better than the other models, with DIC(REP) exp1: 4746.20 and exp2: 9601.48, while the IER model ranked second (DIC(IER) exp1: 5390.61; exp2: 10629.80; see Figure **4B** and Supplementary Table 4 for all model comparisons). Further, we analyzed two datasets from Bavard et al. (2023), which use a Gaussian reward schedule comparable to our tasks g1 and g2, but with the added complexity of a larger context-space (four instead of two choice contexts) and blocking the best option, i.e., making it unavailable, in two out of four choice contexts in a subset of trials. The authors of the original paper concluded that participants employ scaled range normalization rather than divisive normalization. Our analysis shows that the REP model (DIC(REP) Exp3a: 21987.24; exp3b: 24151.11) substantially outperformed the authors’ best model (DIC(RANGEω) exp3a: 28533.3; exp3b: 30205.7) as well as the recently proposed IER model (DIC(IER) Exp3a: 29062.1; exp3b: 30634.4; see Figure **5B** and Supplementary Table 3 for all model comparisons).

## Discussion

We tested mechanisms driving biased choice preferences, which emerge when options learned in stable decision-making contexts are encountered in novel situations, here mixed contexts. To date, it is unclear whether state-dependent normalization (Louie et al. 2011; Louie and Glimcher 2012; Bavard et al. 2021; Bavard and Palminteri 2023), or the consideration of internal reward signals (Molinaro and Collins 2023b) are sufficient to account for such biases. First, using standard analysis, across a range of value-based decision-making tasks (including gains, losses and different types of reward feedback) and 15 datasets, we found evidence that there is a link between repeating context-specific choices during learning and choice preferences in the transfer phase. Even when controlling for reward effects, the relative frequency with which participants chose an option during learning was linked to cross-context choice preferences, post-task valuations, and value uncertainty. Second, using hierarchical Bayesian modeling, we found that the proposed REP model (unbiased RL combined with a repetition bias) outperformed all alternative models in 12 out of 15 datasets and was the second-best model in the remaining three datasets. These results suggest that repetition is a key mechanism in preference formation.

The finding that making a choice increases the likelihood of repeating that choice in the future aligns well with both historical and contemporary theories of decision-making. Over a century ago, Thorndike’s (1911) *Law of Exercise* highlighted the importance of repeated actions in shaping behavior. Consistent with these findings, we observed that higher choice frequency during the learning phase is strongly correlated with biased preferences during the transfer phase, even when expected values were equal on both absolute and relative scales (Figure 3A and B). Notably, if state-dependent normalization were the primary process underlying such biases, we would not expect to observe such strong correlations. Strikingly, we found choice frequency to be related not only to preferences but also to explicit valuations, as participants tended to assign higher values to more frequently chosen stimuli. This observation aligns well with Brehm’s (1956) theory of post-decision changes in valuation. The free-choice paradigm (Brehm 1956; Sharot et al. 2010; Enisman et al. 2021) has been used extensively to investigate the phenomenon that chosen options are overvalued when contrasted to unchosen ones. Critics of this paradigm have argued that the preference for chosen options may be linked to certainty, i.e. participants might prefer a chosen option because they are less uncertain about its true value (Chen and Risen 2010). Our results support both perspectives: increased choice repetition was associated with greater certainty about the underlying value (Figure **4G & H**). Further, more frequently chosen options, while controlling for an effect of reward, were also consistently overvalued compared to those chosen less frequently (Figure **3D**; see Enisman et al. (2021) for a recent meta-analysis on the free choice paradigm).

Our computational modelling approach pursued the three main goals of (i) replication, (ii) testing predictions, and (iii) reanalysis: First, with task variants p1 and p2 we replicated phenomena reported by Klein et al. (2017) and compared models that can in principle explain a biased preference toward options learned in contexts with higher contrast (i.e., for options with context-specific larger relative or normalized value). We found that the proposed REP model showed the lowest DIC value across all alternative models. This finding of the relevance of a repetition bias is supported by previous related results: For example, studies have shown that incorporating choice history into RL models enhances the prediction of future preferences in both monkeys (Lau & Glimcher, 2005) and humans (Seymour et al. 2012; Nebe et al. 2024; Lai and Gershman 2024; Legler et al. 2024; Froelich et al. 2024). Klein et al. (2017) observed modest evidence for a frequency effect using logistic regression. Bavard et al. (2021) incorporated a habitual (choice trace) component inspired by Miller et al. (2019) into one of their models. Although this model nearly matched the performance of their proposed range adaptation model, they found that a habitual component alone was insufficient to account for the large variations in option values in their dataset. Crucially, their approach, while theoretically related to ours, differs in implementation. In our understanding they only update the habitual strength of the chosen option according to Miller et al. (2019). In contrast, in the model proposed here, the repetition biases for both chosen and unchosen options within a context can increase or decrease as previously proposed (Katahira et al. 2017; Palminteri 2023). The usefulness of this updating rule for the unchosen option hints at context-specific, interconnected updating rules of repetition biases, similar to mechanisms observed in reward-driven valuation (Biderman and Shohamy 2021).

Second, to disambiguate between models, we designed several new tasks to explicitly test unique predictions of the REP model. In task p3 the context with a lower value difference (LC context) was repeated more frequently than the context with a higher value difference (HC context; Figure **2C**). For this task, state-dependent normalization (Klein et al. 2017) and the IER model (Molinaro and Collins 2023b) would both predict a preference for the best HC option due to its higher relative value or reduced counterfactual feedback. Contrary to these predictions, participants exhibited a strong preference for the best LC option (A_LC_ = 0.7), which we found is only explained by the REP model. In tasks p4.1 and p4.2, we designed the relative values (contrast of expected values) within each context to be equal (see Figure **1**). Under these conditions, normalization models would predict indifference between options, while the REP model would explain choices by the task-dependent combination of unbiased learning and a repetition bias. In task p4.1, the REP model was favored by quantitative model comparison. This aligns with the finding of our standard analysis that a rather equal choice frequency during learning leads to indifference during transfer (see Figure **2D & K** and Figure**3A**). Surprisingly, task p4.2 was best explained by the REL model utilizing decaying learning rates while the REP model was the second-best model. This was despite the REL model’s theoretical prediction of indifference, which was clearly not the case for participants’ behavior (see Figure **2L**). These results can be explained by two factors. The first factor is that the task was relatively easy to learn (see Figure 2L and methods for details). Therefore, in principle, model comparison will favor models with less complexity. We note that the REL model is the simplest of the models compared, i.e. it has the fewest parameters, relying only on the learning mechanism and a context-dependent difference value ( *Q*_*difference*_). The second factor is the REL model’s parameter estimates (variant with learning decay). We found that the RL model inferred larger Q-value differences between options C_HG_ (0.76) and D_HG_ (0.56) compared to options A_LG_ (0.60) and B_LG_ (0.40). Consequently, the REL model accounts for the observed preference for option C_HG_ over option A_LG_ during the transfer phase, despite its theoretical prediction of indifference. This is further supported by a much higher DIC value of the REL model when employing a constant learning rate (see Supplemental Table 3). These counterintuitive but explainable findings highlight the importance of evaluating models across different learning mechanisms and a diverse range of tasks.

Third, we reanalyzed more complex existing datasets featuring a higher number of choice contexts (see methods for other selection criteria), specifically four different contexts (with eight and twelve different options during learning and 28 and 66 transfer comparisons in contrast to two (probabilistic tasks) or nine (Gaussian tasks) transfer comparisons in our datasets). In these tasks participants encountered scenarios with (i) varying probabilistic gains and losses (Bavard et al. 2018) and (ii) blocked best options in a subset of trials (Bavard and Palminteri 2023). In both of these datasets the REP model performed substantially better than all alternative models (expressed as larger absolute and relative DIC differences when compared to model comparisons on other datasets, see Figure 5B and Supplemental Table 3). In the Bavard et al. (2018) study, the mechanism of the REP model explains the findings of the standard analysis that losses (small or big) or relative losses (small gains) are preferred to larger gains when the options associated with these losses were selected more frequently during learning (see Figure **3C**). This demonstrates a relationship between loss preference and relative choice frequency, which is not implemented in any other model. Similarly, we find by modelling both datasets from Bavard and Palminteri (2023) that middle-valued option that had been repeated more often, due to the blocking of the best option, were preferred more frequently during the transfer phase over their equally valued counterparts (see upper right green rectangle in Figure **3A**).

**Figure 5.**
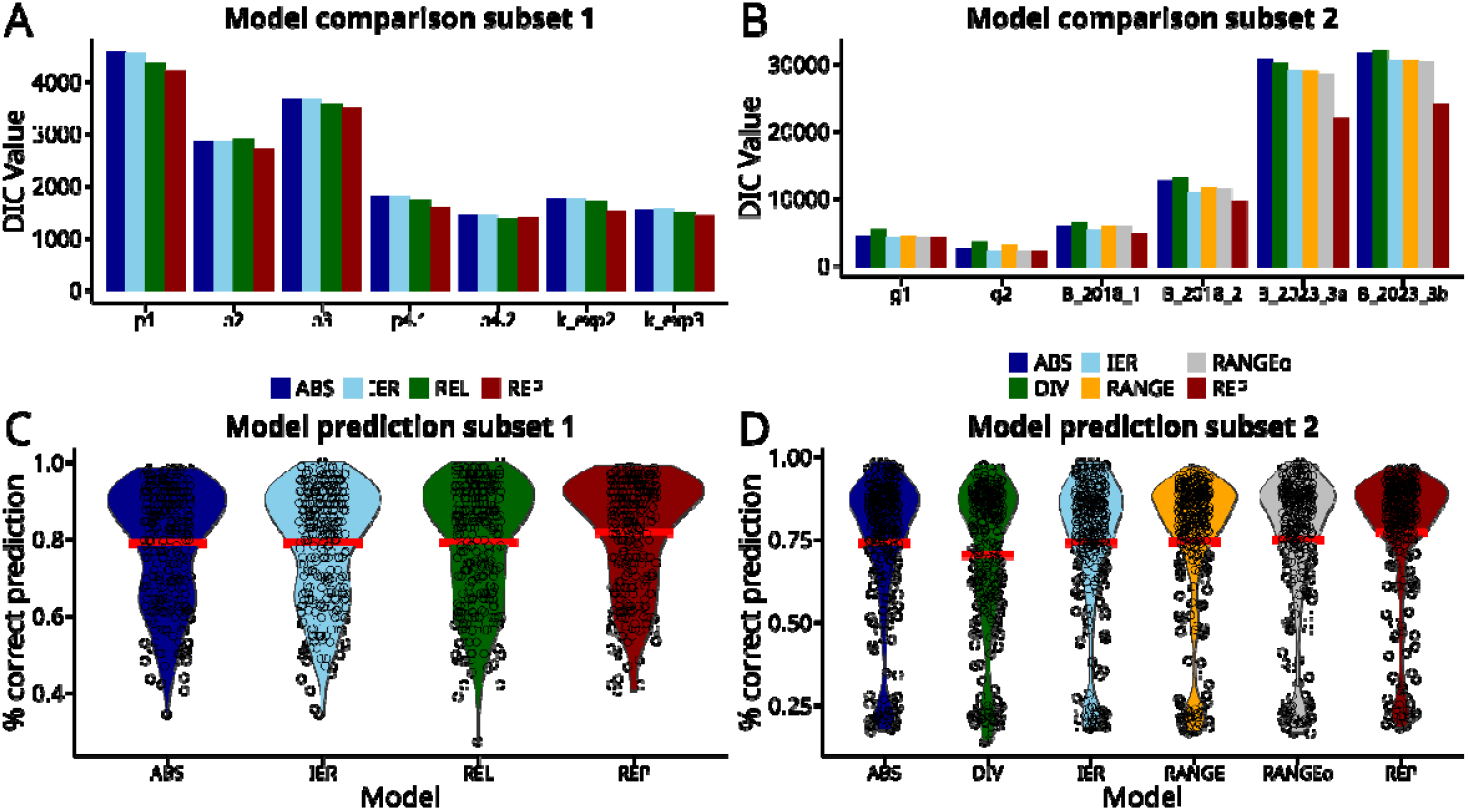
Model comparisons and posterior predictive checks for probabilistic and Gaussian tasks. **A:** Model comparison for probabilistic tasks (p1, p2, p3, p4.1, p4.2) and Experiments 2 and 3 from Klein et al. (2017) based on Deviance Information Criterion (DIC) values, where lower a DIC indicates a better fit. The REP model (red) consistently demonstrates the best fit across all probabilistic tasks except for task 4.2 where it was the second-best model. Specifically, the REP model outperforms all other models in six out of seven tasks or eight out of nine when considering tasks p1.1–p1.3 separately (see Supplemental Table 3). In contrast, other models such as REL (green), IER (blue), and ABS (dark blue) generally perform worse, except for the REL model in task p4.2. **B:** model comparison for Gaussian tasks (g1, g2) and four datasets from Bavard et al. (2018 and 2023). Here, the REP model (red) again is the best model in all cases except for task g1 and g2. Note that for tasks g1 and g2, all models perform approximately equally, likely due to the relative ease of these tasks, where even the ABS model shows nearly similar performance. Other models, including RANGE (yellow), DIV (green), and IER (blue), are generally outperformed by REP, especially in more complex tasks such as those in Bavard et al. (2018, 2023), where participants encountered four different choice contexts during learning and 28 and 66 distinct comparisons during transfer. **C:** posterior predictive checks for all probabilistic tasks, including those from Klein et al. (2017). The plots show the percentage of correct predictions made by each model. The REP model (red) achieves the highest average percentage of correct choice predictions (81.8%), relative to the REL (79.3%), IER (78.9%), and ABS (78.8%) models. Vertical red lines represent the average percentage of correct predictions for each model, while black circles indicate predictions for individual participants. **D:** Gaussian tasks and datasets from Bavard et al. (2018, 2023). Percentage of correct predictions by each model. Consistent with the findings for probabilistic tasks, the REP model (red) performs best in terms of correct predictions (77.2%), whereas models such as RANGEω (75.0%), RANGE (74.5%), IER (74%), ABS (73.7%) and DIV (70.5%) exhibit lower overall performance.

Given these findings, the question arises: why does repetition play such a prominent role in decision-making at all? One explanation is that humans and other animals tend to repeat choices independently of reward to minimize policy complexity and therefore reduce cognitive costs (Gershman 2020; Bhui et al. 2021). A recent study demonstrated that policy compression, i.e. repetition of choices, increases when memory load is high (Lai and Gershman 2024). Similarly, we find that implementing a repetition bias appears to have larger effects on model performance in more complex tasks (see Figure **5B**), i.e., in tasks with more contexts, more transfer comparisons, and more variations in reward magnitude than in simpler tasks where cognitive constraints likely play a lesser role (e.g., g1 and g2). This may not only explain why the REP model was not the best model in tasks g1 and g2 but also suggests that the brain may increasingly rely on specific strategies to manage the heightened cognitive load associated with greater task complexity. One such strategy may be reducing complexity by chunking information into distinct contexts (Heald et al. 2023). By organizing information into contexts, contextual cues in principle enable the brain to rapidly infer the relevant context and activate context-specific relevant memories or action plans (Heald et al. 2021; Cuevas Rivera and Kiebel 2023; Schwöbel et al. 2024). We speculate that contextual inference may be the basis for a context-specific repetition bias, which shapes context-specific representations as recently shown in a modeling study (Schwöbel et al. 2021). Consequently, contextual inference in combination with repetition biases may result in enhanced efficiency of retrieving relevant actions, i.e. effectively speeding up information processing within familiar contexts or when contextual cues are present (Akaishi et al. 2013; Unkelbach and Rom 2017; Braun et al. 2018; Mattavelli et al. 2023; Froelich et al. 2024).

One important advantage of the REP model is that it performs consistently across a range of diverse task parametrizations with varying reward structures, amount of trials per context, and independent of whether partial or full feedback is provided. Crucially, in the REP model, action selection is based solely on variables that are always accessible to the agent, as value-free repetition explicitly does not depend on the outcomes of other options. In contrast the IER (Molinaro and Collins 2023b) and normalization models DIV or RANGE (Louie et al. 2011; Louie et al. 2013; Bavard et al. 2021; Bavard and Palminteri 2023), are at a potential disadvantage here as they work best with full knowledge of the outcome structure, which may or may not be available to participants in both experimental tasks and, importantly, in real-world interactions.

Note we do not state that mechanisms like normalization or intrinsic goal signals are absent in value-based decision-making. Our intention was to explain choice preference by context-specific repetition, a factor not implemented in most alternative models on context-dependent decision biases. However, it is well possible, that multiple mechanisms are at play. For example, the IER model proposed by Molinaro and Collins (2023b) effectively explains switching or exploratory behavior following counterfactual rewards during learning. This is enabled by its explicit mechanism that generates signals based on potentially available counterfactual or non-optimal feedback. Furthermore, other mechanisms are known to influence decision-making under risk (Robinson and Tymula 2019) or when alternative options interact with goals, for instance, anchoring effects in purchasing decisions (Tversky 1972; Tversky and Kahneman 1981; Simonson and Tversky 1992; Furnham and Boo 2011; Daviet and Webb 2023). Future research should examine in more detail if and how repetition and other mechanisms interact, i.e. in multi-attribute decisions in more complex scenarios.

While there is overwhelming consensus on the role of reward in shaping preference and guiding action, i.e. the *Law of Effect* (Thorndike 1911), expected utility theory (Samuelson 1937), operant conditioning (Skinner 2019), prospect theory (Kahneman and Tversky 1979), incentive salience (Robinson and Berridge 1993), the role of repetition in shaping choice preference is still under debate. However, several findings across various domains, i.e. in experimental research on value-based choice (Lau and Glimcher 2005; Seymour et al. 2012; Nebe et al. 2024; Legler et al. 2024; Lai and Gershman 2024; Palminteri 2023; Froelich et al. 2024), perceptual decision-making (Akaishi et al. 2013; Braun et al. 2018), social psychology (Unkelbach and Rom 2017; Mattavelli et al. 2023), recent modelling studies on the interaction between habitual and goal-directed behavior (Miller et al. 2019; Schwöbel et al. 2021), cognitive control (Schwöbel et al. 2024) and real-world preferences (Riefer et al. 2017) point to a fundamental role of the *Law of Exercise* (Thorndike 1911) in shaping human choice. The present study adds to these findings by demonstrating a clear role of context-specific choice repetition in shaping decision biases, providing a foundation for future research to build upon.

## Methods

### Participants

Participants were recruited via the online platform *Prolific* (www.prolific.com). All participants (n = 351; m = 175, w = 176; mean(age) [sd] = 32.32 [8.24] years) were provided written informed consent prior to task participation via the *REDCap* platform. Inclusion criteria were fluent English and age within the range of 18 to 49 years. The study procedure was approved by the Ethics Committee of TU Dresden (ID: 578122019).

### Tasks

All tasks consisted of a learning stage (60 to 80 trials) and a subsequent transfer stage (20 to 36 trials; see Supplemental Table 4). The learning phase of each task consisted of two choice contexts. A choice context was defined by the available options (represented by abstract stimuli). Participants encountered two options per choice context in tasks with probabilistic reward feedback (probabilistic tasks: p1[p.1.1, p1.2, p1.3], p2, p3, p4.1, p4.2) and three options per context in tasks with Gaussian reward feedback (Gaussian tasks: g1, g2). In these probabilistic tasks each option was associated with a reward probability, e.g. 0.7 for winning 0.1€ or nothing. In Gaussian tasks the outcome for each option was sampled from a normal distribution with an option-specific mean, e.g. 70 points, and a standard deviation of 6 (see Supplemental Table 4 for details). Participants received either partial feedback, i.e. they are visually informed about the outcome of the chosen option, or full feedback, i.e. they are informed about the outcome of the chosen option and all alternatives. For a detailed overview of all features for all tasks see Figure 1 and Supplemental Table 4.

### Probabilistic tasks

Task p1 (including variants p1.1, p1.2, and p1.3) and task p2 aimed to replicate the task structure of previous studies. Specifically, p1.1, p1.2, and p1.3 were variants of the task used by Klein et al. (2017), with small variations to the reward probabilities for each option (see Supplemental Table 4 for an overview). In these tasks (p1.1-p1.3), participants received full feedback. Task p2 shared the same underlying parametrization as p1.1 but provided participants with partial feedback. The common feature across these tasks and the Klein et al. (2017) tasks is that the difference in reward probabilities for options in the first context is different from the difference in reward probabilities for options in the second context. For example, during learning, participants encountered a low contrast (LC) context with options A_LC_ (70% reward probability) and B_LC_ (50%), resulting in a 20% difference. Additionally, they faced a high contrast (HC) context with options C_HC_ (70%) and D_HC_ (20%), where the reward probability difference was larger at 50%. By using different task parametrizations and providing either full feedback (p1) or partial feedback (p2), we aimed to replicate the main findings from Klein et al. (2017) and test the robustness of these effects.

In task p3, the LC context was encountered more frequently than the HC context during the learning phase (50 times vs 30 times), allowing us to explicitly test the hypothesis that the number of repetitions influences preferences for choices in the transfer phase.

In tasks p4.1 and p4.2, both contexts in the learning phase had the same difference of reward probabilities (0.2). Participants encountered a low gain (LG) context with options A_LG_(0.60) and B_LG_(0.40), and a high gain (HG) context with options C_HG_(0.76) and D_HG_(0.56), meaning the best options in each context differed in expected value. During the transfer phase, our focus was on testing specific predictions by different models whether participants would prefer option A (0.60) or C (0.76). For example, according to state-dependent normalization, participants should show indifference between these options. In task p4.1, early on in the learning phase, we showed counterfactual feedback to participants in the HG context. This will make the best option in the HG context, C_HG_(0.76), harder to learn and thus less frequently chosen. In task p4.2, this early counterfactual feedback in the HG context was totally absent, making the task easier and we expected that participants chose the best option in the HG context more often. For all probabilistic tasks, reward sequences were pseudorandomized (except the manually altered change in first trials of p4.1 and p4.2), and all participants experienced the same sequence (see Supplemental Table 5 for exact sequences). Crucially, sequences for the learning phase were designed such that the sampled reward probabilities closely matched the true reward probability of each option, measured in blocks of 10 to 12 trials. For instance, if an option had an expected value of 0.7, approximately 7 out of every 10 to 12 trials had to be rewarded. Additionally, the reward sequence for the best options, A_LC_(0.7) and C_HC_(0.7) in both contexts was kept identical for the last four trials in the learning phase (except in task p1.2). This was done to control for simple forms of perseveration or forgetting. For the first transfer trial reward feedback was equal for the best HC and LC options (e.g. A_LC_(0.7) and C_HC_(0.7)).

### Gaussian tasks

Tasks using choice contexts with different reward magnitudes and different reward ranges provide a unique opportunity to test the predictions of normalization models, such as divisive normalization (Louie and Glimcher 2012) and variants of range normalization (Bavard and Palminteri 2023). In tasks g1 (full feedback) and g2 (partial feedback), each choice was associated with a different reward magnitude. Reward magnitudes were drawn from a Gaussian distribution with different means and a standard deviation of six. Participants encountered three options per context. In the LC context, the means of the reward magnitudes were A_LC_(70), B_LC_(60), and C_LC_(50) points, while in the HC context, they were D_HC_(70), E_HC_(40), and F_HC_(10) points. According to standard range normalization participants should show indifference between A_LC_(70) and D_HC_(70), as both would normalize to 1 when applying range normalization. One purpose of using these tasks was to test whether this theoretical prediction holds or if alternative models, especially the proposed model explain the preferences in the transfer phase better. Rewards during the learning phase were sampled from a normal distribution on a trial-by-trial basis (as in Bavard and Palminteri 2023), resulting in unique reward sequences for each participant due to sampling noise. No reward feedback was provided in the transfer phase for Gaussian tasks.

### Bonus payment

For all tasks performance bonuses were computed based on individual task outcomes. Participants could earn an additional 0.1€ for each reward collected in all probabilistic tasks and 0.01 * reward magnitude in points for Gaussian tasks. Bonus reimbursement was paid via the *Prolific* internal payment system after task performance.

### Overall procedure for new tasks

Nine groups of male and female human participants (n = 351) performed nine variants (p1 [p1.1, p.1.2, p1.3], p2, p3, p4.1, p4.2, g1, g2) of value-based decision-making tasks. Each group completed a single task (see task details above). After reading the instructions, participants completed a short training session (10 trials) and then answered four questions, in the form of a quiz, to ensure they understood the task. In case of an error, participants were allowed to retake the quiz until all questions were answered correctly.

Participants made their choices using a computer mouse or touchpad by pressing the left mouse button or touchpad. Keyboard-only input was not allowed, as we assumed participants would be more focused if action selection required using a cursor to indicate their choice, rather than simply pressing left or right keys. In the main task, participants completed a 60 to 80 trial learning phase (depending on the task version, see above), followed by a mandatory three-minute break. Once a counter indicated the end of the break, participants received an on-screen text message informing them that they could resume making choices among all previously encountered options, by pressing the space key. No additional instructions were provided regarding the recombination of options across choice contexts. In the transfer phase, participants made choices between options from different choice contexts. After completing the transfer phase, participants were asked to rate their belief about the true reward probability of each option using a 0-100 visual analogue scale.

### Standard analysis

As a first step, we performed standard analyses based on linear inference statistics. To this end, in the main text we first show descriptive plots of the average rate of choosing the best option during learning and choice preference during the first two transfer trials (TT). To test whether participants show systematic biases in the first TT we conducted exact binomial tests to determine whether preferences between the two high-reward options in the first transfer trial (e.g., A_LC_ [0.7] vs. C_HC_ [0.7]) significantly deviated from chance levels, indicating indifference.

We then focused our analysis on linear relationships between relative choice frequency during learning and relative choice frequency (choice preference) during transfer.

To do this, we counted the choices on the group level for each option during the learning phase and during the transfer phase and normalized these two counts with respect to the total number of presentations of a specific option pair. For example, if option A_LC_ was chosen 400 times and option C_HC_ 600 times, during learning, the relative choice frequency of A_LC_ would be 0.4 and of C_HC_ 0.6 (relative frequency learning). Likewise, if A_LC_ was chosen 25 times and C_HC_ was chosen 75 times when directly compared in the first transfer trial the relative choice frequency would be for A_LC_ 0.25 and for C_HC_ 0.75 (relative frequency transfer). Subsequently, we analyzed the resulting task-specific correlation coefficients between the relative frequency scores during learning and transfer. To further control for potential confounding effects of expected value on the observed correlations, we implemented additional selection criteria for our pairwise comparisons. First, we selected only those pairs where both options had equal expected values (objective expected value), such as A_LC_(0.7) and C_HC_(0.7). Second, to account for possible effects of state-dependent normalization, assuming that participants learn values on a relative scale, we included only pairwise combinations with equal relative value. Additionally, we excluded the worst options (14-point bandits) from Bavard and Palminteri (2023), as these options were selected in less than 10% of trials within a three-option context on average. However, we also computed all correlations with these options included, and the results did not substantially change (see Supplemental Table 2). For the analysis of the relationship between choice frequency and valuation we determined the relative choice frequency for each option at the individual participant level. For each option, we calculated a normalized frequency score, representing how often it was chosen relative to all other options encountered in the task. We then determined the differences in these normalized choice frequencies between pairs of options and correlated these differences with the differences in their individual valuation ratings. To control for reward effects, we focused on options with equal expected values (absolute and relative), as described previously. To analyze the relationship between choice frequency and uncertainty, we calculated the relative choice frequency for each option and the standard deviation of valuation ratings at the group level. We then tested for a linear relationship between relative choice frequency and the standard deviation of the value distribution. We further examined a correlation of relative choice frequency and absolute distance from the true reward probability on the participant level (probabilistic tasks) or reward magnitude (Gaussian tasks). Data normality was assessed using the Shapiro-Wilk test. Based on test results, we report either Pearson or Spearman correlation coefficients.

### Previously published datasets

We reanalyzed six datasets from three previous studies (Klein et al. 2017; Bavard et al. 2018; Bavard and Palminteri 2023). The selection of datasets was made prior to conducting any analyses (see Table 1 for details.).

**Table 1.** Reanalysis of six previously published datasets.

| **Paper** | **Tasks** | **Reason for Reanalysis** |
| --- | --- | --- |
| **Klein et al. 2017** | Exp2 (n = 24), Exp3 (n = 21) | Experiments 2 and 3 support the authors main findings, i.e. these tasks provide evidence for relative value learning. We replicated their task design in our tasks p1 and p2. |
| **Bavard et al. 2018** | Exp1 (n = 20), Exp2 (n = 40) | This study investigated choices between losses and gains and gains with different magnitude in the transfer phase. Further this dataset is of higher complexity due to its four choice contexts and 28 distinct transfer comparisons in contrast to only one transfer comparison in Exp.2 and Exp.3 by Klein et al. (2017) and two transfer comparisons in our tasks p1 and p2. The observed apparent preference for losses is modelled by state-dependent normalization. |
| **Bavard & Palminteri 2023** | Exp3a (n = 100), Exp3b (n = 100) | This study established evidence that preferences in the transfer phase are best explained by scaled range normalization (denoted as RANGEω). In addition, Bavard and Palminteri (2023) blocked the best option in a subset of contexts during learning. This enforces repetition of the second-best option. Our Gaussian tasks (g1 and g2) are similar to their experiments 3a and 3b in terms of reward feedback but of lower complexity (4 transfer comparisons in our datasets vs. 66 transfer comparisons in their dataset) |

### Reinforcement Learning (RL) models

Multiple computational models have been proposed to explain decision biases, i.e. irrational choice preferences, in value-based decision-making (Louie et al. 2013; Klein et al. 2017; Bavard et al. 2018; Bavard and Palminteri 2023; Molinaro and Collins 2023b). In the present study we systematically compare these models against each other. Specifically, we analyzed our data using six different RL models: relative value learning (Klein et al. 2017), divisive normalization (Louie et al. 2011; Louie and Glimcher 2012; Webb et al. 2021), variants of range normalization (Bavard & Palminteri, 2023b) and a model that weights external and internal reward signals (Molinaro and Collins 2023a). As a control model, we used an absolute value learning (ABS) model (Sutton and Barto 2018). In the ABS model, which is equivalent to standard Q-learning, Q-values within a context are updated in a stepwise procedure through a prediction error term:

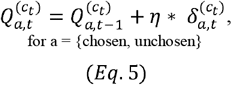

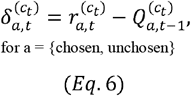

where *c* is the context in trial *t. η* is a placeholder for the learning-rate (for different parametrizations see below) and *r*_*a*_ is the reward for the chosen or unchosen option in trial *t*. *δ*_*a*,*t*_ is the corresponding prediction error, i.e. the difference between the received reward *r*_*a*,*t*_ and the previously estimated Q-value for that option 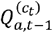. In the case of partial feedback, only the Q-value for the chosen option is updated, thus *a* = chosen. In the case of full feedback, Q-values for both the chosen and unchosen options are updated and *a* = {chosen, unchosen}. Across all probabilistic tasks and the data from Klein et al. 2017 participants encountered one unchosen option per context. In Gaussian tasks and data from Bavard and Palminteri 2023 participants encountered two unchosen options per context.

### Learning rules

Before presenting the remaining models, and to finalize the ABS model, we introduce two learning mechanism variants, which were across all models.

In the first model variant we used a trial-wise learning rate that decays as a function of context occurrences and in the second model variant we used different constant learning rates for chosen and unchosen options. Model variant 1 is motivated by RL research and suggests that when reward probabilities are stationary (no fluctuations in the environment), learning rates are assumed to decay over time, i.e. participants go gradually from a phase of initial exploration to exploitation (Sutton & Barto, 2018). A constant learning rate throughout the entire experiment would not adequately capture this shift. We tested several decay functions through comparative analyses based on simulated data (details not reported). Based on these tests, we selected a hyperbolic decay function, where the learning rate decreases as a function of the number of times a context has occurred (Eq. 9).

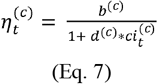

Here *b*^*(c)*^ is a context-specific baseline learning rate at the beginning of the learning phase, is the context-specific decay parameter that weights the number of times a specific context has occurred and 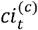 is a context-specific counter that increases every time the corresponding context *c is* experienced. 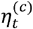is the corresponding context and trial-specific learning rate.

For the second model variant, following prior studies, we implemented different constant learning rates for chosen (*δ* _*chosen*_) and unchosen (*δ*_*unchosen*_) options (Klein et al. 2017; Bavard et al. 2018; Bavard and Palminteri 2023). Note we do not systematically compare these two model variants, as this would go beyond the scope of this study. Rather, we implemented both learning rule variants to show that our main results do not change when other learning mechanisms, especially those that were used in prominent studies on state dependent normalization are applied (Palminteri et al. 2015; Bavard et al. 2018; Bavard and Palminteri 2023).

### Repetition model

The proposed repetition model (REP) consists of two components: (i) standard reinforcement learning (ABS; see Eq. 5 and Eq.6 above) with (ii) reward-independent learning of a bias to repeat previous choices (Akaishi et al. 2013; Katahira et al. 2017; Schwöbel et al. 2021; Palminteri 2023; Schwöbel et al. 2024). For this second component we let repeated choices influence the repetition bias (REPbias) of an option depending on which option is chosen:

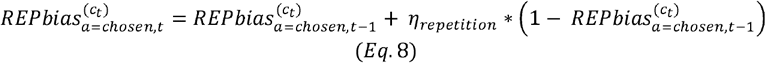

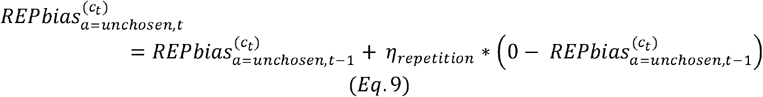

i.e., after each choice, the REPbias within the current context increases for the chosen option and decreases for the unchosen options. The trial-specific change in an option’s REPbias is governed by a participant-specific learning rate parameter, *η*_*repetition*_. This model is theoretically related to Miller et al. (2019; Bavard et al. 2021) but differs in two significant ways: Unlike Miller et al., we do not employ a weighting parameter (arbiter) to balance the influence of Q-values and REPbiases. Further, following (Katahira et al. 2017; Palminteri 2023), the REP model allows the REPbias of each option to both increase and decrease, providing a more flexible adjustment based on the choice history of each participant. This contrasts with models that may only allow for the strengthening of biases over time.

### Divisive normalization

The idea behind divisive normalization (DIV) applied to value-based decision making is that rewards are normalized with respect to other rewards within a context (Louie et al. 2011, Louie and Glimcher 2012):

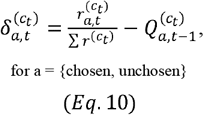

particularly both the chosen and unchosen (if full feedback is provided) rewards *r*_*a*,*t*_ within the current context *c*_*t*_ are normalized relative to the sum 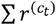of alternative rewards within the prediction error term 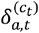. Q-values are updated as usually according to Eq. 5 but with the modified prediction error term (Eq. 10).

### Range normalization

Range normalization (RANGE) assumes that option values are rescaled as a function of the maximum and the minimum rewards presented in a context (Bavard et al. 2021):

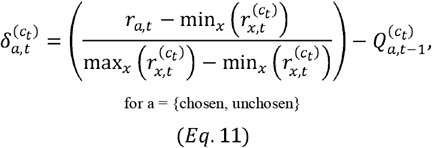

here, the chosen and unchosen (in case of full feedback) reward *r*_*a*,*t*_ is normalized as a function of the minimum (worst) and maximum (best) reward in the current context *c*_*t*_ (Eq. 11). This normalized reward within the prediction error term is then used to update the Q-values for the corresponding options (Eq. 5).

### Scaled range normalization

Bavard and Palminteri (2023) proposed an extension to range normalization by introducing anω exponent enhancing the flexibility of the normalization process:

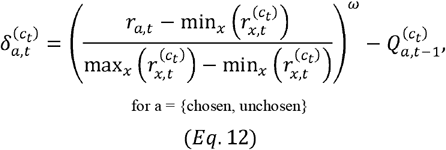

In this scaled range normalization (*RANGE*) model, the ω parameter allows the model to skew value estimates either concavely or convexly (Eq. 12). This skewing was argued to better account for the non-linearity in choice preferences observed in the datasets analyzed by Bavard and Palminteri (2023).

### Intrinsic enhanced reward model (IER)

Molinaro and Collins (2023b) proposed a goal-centric RL model that integrates an intrinsic reward mechanism. This binary internal signal (0 or 1) corresponds to the goal of selecting the best option in each trial. The model weights this intrinsic signal with external reward feedback, allowing Q-values to be updated as a weighted combination of both outcomes. The underlying idea is that participants prefer options during the transfer phase that are more consistent with their internal goal of choosing the best option:

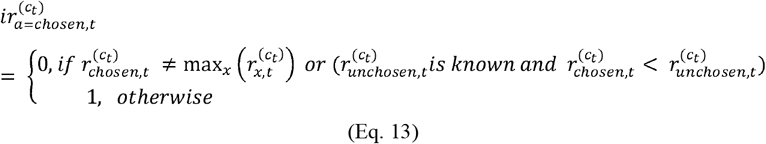

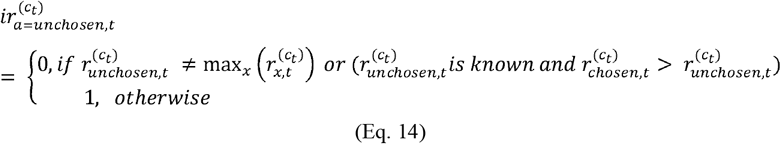

In words, the trial-wise intrinsic reward signal 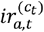for an option is set to 0 if (i) the chosen reward does not equal the maximum obtainable reward in that context or if (ii) the chosen reward turns out to be smaller than the unchosen reward. Otherwise, 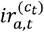is set to 1. The intrinsic reward signal in that context is then weighted relative to the extrinsic reward *r*_*a*,*t*_:

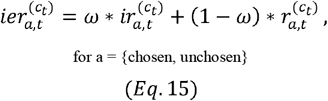

where*ω* is a weighting parameter. Subsequently, Q-values are updated according to Eq. 5, but using prediction errors 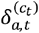based on the intrinsically enhanced reward 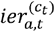signal:

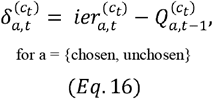

### Relative model (REL)

In the relative value learning (REL) model proposed by Klein et al. (2017), participants are assumed to directly learn a *Q*_*difference*_ value for each context *c*:

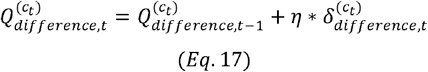

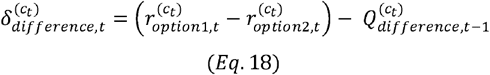

where reward information for both options within the current context are used to compute prediction error and update the *Q*_*difference*_ estimates. This means that the REL model can only apply to choice contexts with two options.

### Action selection

Action selection for each model was computed using a softmax decision rule (Sutton & Barto, 1998), which weights Q-values (Eq.19; ABS, DIV, RANGE, RANGE IER *ω* models), _*Qdifference*_ values (Eq.21; REL model) or Q-values and REPbiases in the case of the REP model (Eq.20), to model the probability of selecting the chosen option on trial *t*.

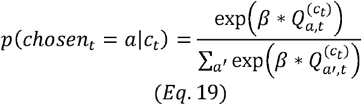

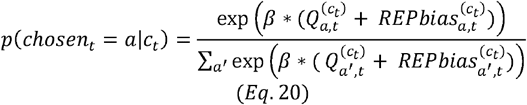

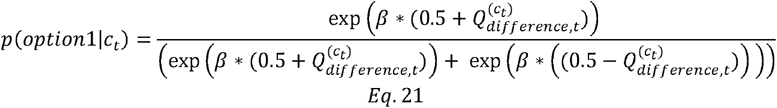

All three softmax functions are parameterized by the inverse temperature *β*. When *β*= 0, choices are random, and as *β*increases, choices become more dependent on learned Q-values.

For probabilistic tasks, following Klein et al. (2017) and as in the standard analysis (see above) we focused our analysis on the first two transfer trials for each choice context. Previous studies have similarly focused on the initial decisions in the transfer phase or included only two or four decisions per transfer pair (Klein et al., 2017; Bavard et al., 2018; Bavard & Palminteri, 2023). For Gaussian tasks we included all four transfer trials for each context. For our new probabilistic tasks learning mechanisms were applied to the learning and transfer phases, as feedback was provided during transfer. The reptition-bias mechanism was applied during learning and transfer for all tasks. Note results did not differ to the case when the repetition bias was only applied to the learning phase (see Supplemental Table 3).

### Modeling partial feedback

In the partial feedback condition, reward information for alternative (unchosen) options, such as their outcomes (rewards) or the range of possible alternative rewards, was unavailable to participants. This limitation presents a challenge for accurately updating Q-values and applying normalization or weighting in the DIV, RANGE, RANGEω, and IER models. Past studies have addressed this issue in different ways. Some researchers treated the learned and stored Q-values as valid proxies for the unobserved rewards. These proxy values were then used as inputs for normalizing within the prediction error terms (Palminteri et al., 2015; Bavard et al., 2018). Others opted for computational simplicity by allowing models to retain full knowledge of contextual information, such as the minimum and maximum possible rewards, even though participants did not receive complete feedback (Molinaro & Collins, 2023b). For comparing these models in our study, we addressed the challenge of partial feedback for these models by implementing a memory system for the DIV, REL, RANGE, RANGEω, and IER models. When participants received partial feedback, the most recent known reward information for each option was retrieved to inform their computations. Additionally, as proposed by Molinaro and Collins (2023b), we calculated the intrinsically enhanced reward (IER) signal exclusively for the chosen option when full feedback was unavailable. Note that the REP model computations are in all conditions, including partial feedback, only based on information that is always available to the agent, i.e. in Eqs. 5, 6, 8 and 9, the agent only updates context-specific values, depending on the selected action.

### Hierarchical Bayesian inference

Models were fit to all trials from all participants using a hierarchical Bayesian modeling approach. Parameter inference was performed using Markov Chain Monte Carlo sampling as implemented in the JAGS software package (Plummer, 2003) (Version 4.3) in combination with R (Version 4.3) and the R2Jags package. For group-level means, we used uniform priors defined over numerically plausible parameter ranges ([-5, 5] for the learning- (η) and baseline learning-rates *d*^(c)^ in standard-normal space; [0, 5] for *β* and [0,3] for the hyperbolic decay*b*^*(c)*^. Group-level standard deviations were likewise modelled via uniform priors over a plausible parameter range. For full model code and parametrization see data availability section (made public upon publication). We initially ran two chains with a varying burn-in period of at least 20.000 samples and thinning of two. Chain convergence was assessed via the Gelman-Rubinstein convergence diagnostic 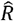and sampling was continued until 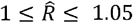 for all group-level and individual-subject parameters. In some cases, learning-rate parameters did exceed that threshold, however all parameters had a 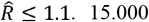 additional samples were retained for further analysis.

### Model comparison

The ABS, REP and IER models were compared across all tasks. For tasks with two options per context and equal reward magnitude (p1, p2, p3, p4.1, p4.2 and data from Klein et al. 2017) we additionally used the REL model. For tasks with more than two options per context and/or where the range of reward magnitudes differed between contexts or options (tasks g1, g2, and datasets from Bavard et al., 2018, 2023), we additionally applied divisive normalization (DIV) (Louie et al. 2011; Louie et al. 2013) and range normalization models (RANGE & RANGE *ω*) (Bavard et al. 2021; Bavard and Palminteri 2023). Quantitative model comparison was performed via the deviance information criterion (DIC) where lower values indicate better fit (Spiegelhalter et al. 2012).

### Posterior predictive checks

We carried out posterior predictive checks to examine whether each model can reproduce participants’ choices in both learning and transfer phase. For validation, we plot for each participant the correct predicted binary choices averaged across 10k datasets simulated from the posterior distributions of all models.

## Code and Data Availability

All model code used to generate the results, and all newly collected datasets will be made public on OSF upon publication.

## Acknowledgements

We thank Yaning Hu for assistance with task programming and Dario Cuevas-Riva for helpful discussions and advice.

## Author Contributions

BJW and SJK conceived the idea. BJW designed the study. BJW and HBW acquired the data. BJW contributed analytical tools. BJW and HBW analyzed the data. BJW wrote the paper. SJK provided funding and supervised the project. All authors read and edited versions of the manuscript and approved the final version.

## Publication bibliography

Akaishi, Rei; Umeda, Kazumasa; Nagase, Asako; Sakai, Katsuyuki (2013): Autonomous Mechanism of Internal Choice Estimate Underlies Decision Inertia. In Neuron 81 (1), pp. 195–206. DOI: 10.1016/j.neuron.2013.10.018.

Bavard, Sophie; Lebreton, Maël; Khamassi, Mehdi; Coricelli, Giorgio; Palminteri, Stefano (2018): Reference-point centering and range-adaptation enhance human reinforcement learning at the cost of irrational preferences. In Nat Commun 9 (1), p. 4503. DOI: 10.1038/s41467-018-06781-2.

Bavard, Sophie; Palminteri, Stefano (2023): The functional form of value normalization in human reinforcement learning. In eLife 12. DOI: 10.7554/eLife.83891.

Bavard, Sophie; Rustichini, Aldo; Palminteri, Stefano (2021): Two sides of the same coin: Beneficial and detrimental consequences of range adaptation in human reinforcement learning. In Science advances 7 (14). DOI: 10.1126/sciadv.abe0340.

Bhui, Rahul; Lai, Lucy; Gershman, Samuel J. (2021): Resource-rational decision making. In Current Opinion in Behavioral Sciences 41, pp. 15–21. DOI: 10.1016/j.cobeha.2021.02.015.

Biderman, Natalie; Shohamy, Daphna (2021): Memory and decision making interact to shape the value of unchosen options. In Nature Communications 12 (1), p. 4648. DOI: 10.1038/s41467-021-24907-x.

Bouton, Mark E.; Maren, Stephen; McNally, Gavan P. (2021): BEHAVIORAL AND NEUROBIOLOGICAL MECHANISMS OF PAVLOVIAN AND INSTRUMENTAL EXTINCTION LEARNING. In Physiological reviews 101 (2), pp. 611–681. DOI: 10.1152/physrev.00016.2020.

Braun, Anke; Urai, Anne E.; Donner, Tobias H. (2018): Adaptive History Biases Result from Confidence-Weighted Accumulation of past Choices. In J. Neurosci. 38 (10), pp. 2418–2429. DOI: 10.1523/JNEUROSCI.2189-17.2017.

Brehm, J. W. (1956): Postdecision changes in the desirability of alternatives. In Journal of abnormal psychology 52 (3), pp. 384–389. DOI: 10.1037/h0041006.

Chen, M. Keith; Risen, Jane L. (2010): How choice affects and reflects preferences: Revisiting the free-choice paradigm. In Journal of Personality and Social Psychology 99 (4), pp. 573–594. DOI: 10.1037/a0020217.

Courville, Aaron C.; Daw, Nathaniel D.; Touretzky, David S. (2006): Bayesian theories of conditioning in a changing world. In Trends in Cognitive Sciences 10 (7), pp. 294–300. DOI: 10.1016/j.tics.2006.05.004.

Cuevas Rivera, Dario; Kiebel, Stefan (2023): The effects of probabilistic context inference on motor adaptation. In PloS one 18 (7), e0286749. DOI: 10.1371/journal.pone.0286749.

Daviet, Remi; Webb, Ryan(2023): A test of attribute normalization via a double decoy effect. In Journal of Mathematical Psychology 113, p. 102741. DOI: 10.1016/j.jmp.2022.102741.

Enisman, Maya; Shpitzer, Hila; Kleiman, Tali (2021): Choice changes preferences, not merely reflects them: A meta-analysis of the artifact-free free-choice paradigm. In Journal of Personality and Social Psychology 120 (1), pp. 16–29. DOI: 10.1037/pspa0000263.

Froelich, Sascha; Wagner, Ben Jo; Smolka, Michael N.; Kiebel, Stefan J. (2024): Context-Dependent Interaction Between Goal-Directed and Habitual Control Under Time Pressure. In bioRxiv, 2024.09.28.615575. DOI: 10.1101/2024.09.28.615575.

Furnham, Adrian; Boo, Hua Chu (2011): A literature review of the anchoring effect. In The Journal of Socio-Economics 40 (1), pp. 35–42. DOI: 10.1016/j.socec.2010.10.008.

Gershman, Samuel J. (2017): Context-dependent learning and causal structure. In Psychonomic bulletin & review 24 (2), pp. 557–565. DOI: 10.3758/s13423-016-1110-x.

Gershman, Samuel J. (2020): Origin of perseveration in the trade-off between reward and complexity. In Cognition 204, p. 104394. DOI: 10.1016/j.cognition.2020.104394.

Gershman, Samuel J.; Blei, David M.; Niv, Yael (2010): Context, learning, and extinction. In Psychological Review 117 (1), pp. 197–209. DOI: 10.1037/a0017808.

Glimcher, Paul W. (2022): Efficiently irrational: deciphering the riddle of human choice. In Trends in Cognitive Sciences 26 (8), pp. 669–687. DOI: 10.1016/j.tics.2022.04.007.

Heald, James B.; Lengyel, Máté; Wolpert, Daniel M. (2021): Contextual inference underlies the learning of sensorimotor repertoires. In Nature 600 (7889), pp. 489–493. DOI: 10.1038/s41586-021-04129-3.

Heald, James B.; Lengyel, Máté; Wolpert, Daniel M. (2023): Contextual inference in learning and memory. In Trends in Cognitive Sciences 27 (1), pp. 43–64. DOI: 10.1016/j.tics.2022.10.004.

Kahneman, Daniel; Tversky, Amos (1979): Prospect Theory: An Analysis of Decision under Risk. In Econometrica 47 (2), p. 263. DOI: 10.2307/1914185.

Katahira, Kentaro; Yuki, Shoko; Okanoya, Kazuo (2017): Model-based estimation of subjective values using choice tasks with probabilistic feedback. In Journal of Mathematical Psychology 79, pp. 29–43. DOI: 10.1016/j.jmp.2017.05.005.

Klein, Tilmann A.; Ullsperger, Markus; Jocham, Gerhard (2017): Learning relative values in the striatum induces violations of normative decision making. In Nature Communications 8 (1), p. 16033. DOI: 10.1038/ncomms16033.

Lai, Lucy; Gershman, Samuel J. (2024): Human decision making balances reward maximization and policy compression. In PLOS Computational Biology 20 (4), e1012057. DOI: 10.1371/journal.pcbi.1012057.

Lau, Brian; Glimcher, Paul W. (2005): Dynamic response-by-response models of matching behavior in rhesus monkeys. In Journal of the Experimental Analysis of Behavior 84 (3), pp. 555–579. DOI: 10.1901/jeab.2005.110-04.

Legler, Eric; Rivera, Darío Cuevas; Schwöbel, Sarah; Wagner, Ben J.; Kiebel, Stefan (2024): Cognitive Computational Model Reveals Repetition Bias in a Sequential Decision-Making Task. In bioRxiv, 2024.05.30.596605. DOI: 10.1101/2024.05.30.596605.

Louie, Kenway; Glimcher, Paul W. (2012): Efficient coding and the neural representation of value. In Annals of the New York Academy of Sciences 1251 (1), pp. 13–32. DOI: 10.1111/j.1749-6632.2012.06496.x.

Louie, Kenway; Glimcher, Paul W.; Webb, Ryan (2015): Adaptive neural coding: from biological to behavioral decision-making. In Current Opinion in Behavioral Sciences 5, pp. 91–99. DOI: 10.1016/j.cobeha.2015.08.008.

Louie, Kenway; Grattan, Lauren E.; Glimcher, Paul W. (2011): Reward value-based gain control: divisive normalization in parietal cortex. In J. Neurosci. 31 (29), pp. 10627–10639. DOI: 10.1523/JNEUROSCI.1237-11.2011.

Louie, Kenway; Khaw, Mel W.; Glimcher, Paul W. (2013): Normalization is a general neural mechanism for context-dependent decision making. In Proceedings of the National Academy of Sciences 110 (15), pp. 6139–6144. DOI: 10.1073/pnas.1217854110.

Mattavelli, Simone; Corneille, Olivier; Unkelbach, Christian (2023): Truth by repetition … Without repetition: Testing the effect of instructed repetition on truth judgments. In Journal of experimental psychology. Learning, memory, and cognition 49 (8), pp. 1264–1279. DOI: 10.1037/xlm0001170.

Miller, Kevin J.; Shenhav, Amitai; Ludvig, Elliot A. (2019): Habits without values. In Psychological review 126 (2), pp. 292–311. DOI: 10.1037/rev0000120.

Molinaro, Gaia; Collins, Anne G. E. (2023a): A goal-centric outlook on learning. In Trends in Cognitive Sciences 27 (12), pp. 1150–1164. DOI: 10.1016/j.tics.2023.08.011.

Molinaro, Gaia; Collins, Anne G. E. (2023b): Intrinsic rewards explain context-sensitive valuation in reinforcement learning. In PLOS Biology 21 (7), e3002201. DOI: 10.1371/journal.pbio.3002201.

Nebe, Stephan; Kretzschmar, André; Brandt, Maike C.; Tobler, Philippe N. (2024): Characterizing Human Habits in the Lab. In Collabra: Psychology 10 (1). DOI: 10.1525/collabra.92949.

Niv, Yael (2019): Learning task-state representations. In Nature Neuroscience 22 (10), pp. 1544–1553. DOI: 10.1038/s41593-019-0470-8.

Palminteri, Stefano (2023): Choice-confirmation bias and gradual perseveration in human reinforcement learning. In Behavioral neuroscience 137 (1), pp. 78–88. DOI: 10.1037/bne0000541.

Palminteri, Stefano; Khamassi, Mehdi; Joffily, Mateus; Coricelli, Giorgio (2015): Contextual modulation of value signals in reward and punishment learning. In Nature Communications 6, p. 8096. DOI: 10.1038/ncomms9096.

Rabinowitz, Neil C.; Willmore, Ben D. B.; Schnupp, Jan W. H.; King, Andrew J. (2011): Contrast gain control in auditory cortex. In Neuron 70 (6), pp. 1178–1191. DOI: 10.1016/j.neuron.2011.04.030.

Riefer, Peter S.; Prior, Rosie; Blair, Nicholas; Pavey, Giles; Love, Bradley C. (2017): Coherency Maximizing Exploration in the Supermarket. In Nature Human Behaviour 1. DOI: 10.1038/s41562-016-0017.

Robinson, Terry; Berridge, Kent (1993): The neural basis of drug craving: An incentive-sensitization theory of addiction. In Brain Research Reviews 18 (3), pp. 247–291. DOI: 10.1016/0165-0173(93)90013-P.

Robinson, Terry; Tymula, Agnieszka (2019): Divisive Normalisation of Value Explains Choice-Reversals in Decision-Making Under Risk. In SSRN Journal. DOI: 10.2139/ssrn.3492823.

Rosas, Juan M.; Todd, Travis P.; Bouton, Mark E. (2013): Context change and associative learning. In Wiley Interdisciplinary Reviews: Cognitive Science 4 (3), pp. 237–244. DOI: 10.1002/wcs.1225.

Samuelson, Paul A. (1937): A Note on Measurement of Utility. In The Review of Economic Studies 4 (2), p. 155. DOI: 10.2307/2967612.

Schwöbel, Sarah; Marković, Dimitrije; Smolka, Michael N.; Kiebel, Stefan (2024): Joint modeling of choices and reaction times based on Bayesian contextual behavioral control. In PLOS Computational Biology 20 (7), e1012228. DOI: 10.1371/journal.pcbi.1012228.

Schwöbel, Sarah; Marković, Dimitrije; Smolka, Michael N.; Kiebel, Stefan J. (2021): Balancing control: A Bayesian interpretation of habitual and goal-directed behavior. In 0022-2496 100, p. 102472. DOI: 10.1016/j.jmp.2020.102472.

Seymour, Ben; Daw, Nathaniel D.; Roiser, Jonathan P.; Dayan, Peter; Dolan, Ray (2012): Serotonin selectively modulates reward value in human decision-making. In J. Neurosci. 32 (17), pp. 5833–5842. DOI: 10.1523/JNEUROSCI.0053-12.2012.

Shapley, R. M.; Victor, J. D. (1978): The effect of contrast on the transfer properties of cat retinal ganglion cells. In The Journal of Physiology 285 (1), pp. 275–298. DOI: 10.1113/jphysiol.1978.sp012571.

Sharot, Tali; Martino, Benedetto de; Dolan, Raymond J. (2009): How choice reveals and shapes expected hedonic outcome. In The Journal of neuroscience : the official journal of the Society for Neuroscience 29 (12), pp. 3760–3765. DOI: 10.1523/JNEUROSCI.4972-08.2009.

Sharot, Tali; Velasquez, Cristina M.; Dolan, Raymond J. (2010): Do decisions shape preference? Evidence from blind choice. In Psychological Science 21 (9), pp. 1231–1235. DOI: 10.1177/0956797610379235.

Simonson, Itamar; Tversky, Amos (1992): Choice in Context: Tradeoff Contrast and Extremeness Aversion. In Journal of Marketing Research 29 (3), pp. 281–295. DOI: 10.1177/002224379202900301.

Skinner, B. F. (2019): The Behavior of Organisms. An Experimental Analysis: B. F. Skinner Foundation.

Spiegelhalter, David J.; Best, Nicola G.; Carlin, Bradley P.; Linde, Angelika (2014): The Deviance Information Criterion: 12 Years on. In J. R. Stat. Soc. Ser. B. Stat. Methodol. 76 (3), pp. 485–493. DOI: 10.1111/rssb.12062.

Sutton, Richard S.; Barto, Andrew G. (2018): Reinforcement Learning, second edition. An Introduction: MIT Press.

Thorndike, Edward L. (1911): Animal intelligence: Experimental studies. In (No Title). DOI: 10.5962/bhl.title.55072.

Tversky, A.; Kahneman, D. (1981): The framing of decisions and the psychology of choice. In Science 211 (4481), pp. 453–458. DOI: 10.1126/science.7455683.

Tversky, Amos (1972): Elimination by aspects: A theory of choice. In Psychological Review 79 (4), pp. 281–299. DOI: 10.1037/h0032955.

Unkelbach, Christian; Rom, Sarah C. (2017): A referential theory of the repetition-induced truth effect. In Cognition 160, pp. 110–126. DOI: 10.1016/j.cognition.2016.12.016.

Webb, Ryan; Glimcher, Paul W.; Louie, Kenway (2021): The Normalization of Consumer Valuations: Context-Dependent Preferences from Neurobiological Constraints. In Management Science 67 (1), pp. 93–125. DOI: 10.1287/mnsc.2019.3536.

Wood, Wendy; Rünger, Dennis (2016): Psychology of Habit. In Annual Review of Psychology 67, pp. 289–314. DOI: 10.1146/annurev-psych-122414-033417.

