## Supplement for "Explaining decision biases through context-dependent repetition"

**Supplemental**

Supplemental Table 1 Model comparison overview. For tasks p1 to p4.2 relative learning and other normalization procedures do not differ in their predictions. Thus, for these tasks relative value learning is the only normalization model. For tasks with different reward magnitudes and/or more than two options per context we instead fitted divisive- and range normalization models.

| Model | Dataset (task) | | | | | |
| --- | --- | --- | --- | --- | --- | --- |
|  | p1.1, p1.2, p1.3, p2, p3, p4.1, p4.2 | g1, g2 | Exp2, Exp 3  (Klein et al. 2017) | | Exp1, Exp2  Bavard et al. 2018 | Exp3a, Exp3b  (Bavard and Palminteri 2023) |
| REP (standard Q-learning  + repetition bias) | x | x | x | x | | x |
| ABS (standard Q-learning) | x | x | x | x | | x |
| IER (intrinsic enhanced reward; Mollinaro & Collins 2023b) | x | x | x | x | | x |
| REL (relative Q-learning; Klein et al. 2017) | x | - | x | - | | - |
| DIV (divisive normalization (Louie and Glimcher 2012) | - | x | x | x | | x |
| RAN (range normalization; Bavard and Palminteri 2023) | - | x | x | x | | x |
| RAN $\boldsymbol{\omega}$ (range normalization + $\boldsymbol{\omega}$ exponent ; Bavard and Palminteri 2023) | - | x | x | x | | x |

Supplemental Table 2 Correlation coefficients for relative choice-frequency correlation between learning and transfer phase for all options pairs encountered during transfer. For tasks p1, p2, p3, p4.1, p4.2 (first row) and g1, g2 (second row) data was pooled across tasks.

| Pearson R | | Spearman R | | p-value  (Pearson) | | p-value  (Spearman) | | Task | | Shapiro-  Wilk  (learning) | | | Shapiro-  Wilk  (transfer) |
| --- | --- | --- | --- | --- | --- | --- | --- | --- | --- | --- | --- | --- | --- |
| 0.80 | | 0.82 | | 2.01e-04* | | 1.30e-04* | | p1-p4.2 | | | 1.32e-01 | | 1.09e-01 |
| 0.94 | | 0.98 | | 4.14e-09* | | 5.40e-13* | | g1-g2 | | | 5.94e-02 | | 2.81e-03* |
| 0.72 | | 0.73 | | 1.62e-05* | | 1.01e-05* | | B2018_1 | | | 3.06e-01 | | 1.16e-01 |
| 0.91 | | 0.83 | | 1.99e-11* | | 3.51e-08* | | B2018_2 | | | 8.40e-03* | | 5.37e-02 |
| 0.92 | | 0.92 | | 5.86e-29* | | 4.35e-27* | | B2023_1 | | | 1.70e-04* | | 1.36e-05* |
| 0.91 | | 0.86 | | 1.49e-25* | | 8.41e-21* | | B2023_2 | | | 8.76e-05* | | 5.60e-05* |
| 0.88 | | 0.87 | | 8.27e-73* | | 1.57e-70* | | all datasets | | | 1.50e-05* | | 6.26e-09* |
| 0.54 | | 0.42 | | 5.79e-04 | | 8.81e-03 | | equal absolute value (all comparisons from all datasets) | | | 8.53e-04* | | 2.49e-02 |
| 0.88 | | 0.80 | | 2.55e-07* | | 1.94e-05* | | loss preference | | | 1.36e-02 | | 3.40e-01 |
| 0.60 | | 0.66 | | 1.42e-03* | | 3.27e-04* | | equal absolute value (without low value (14 points) bandits from B2023) | | | 2.09e-03* | | 1.54e-03* |
| 0.71 | | 0.67 | | 0.00087* | | 0.002181* | | Equal relative values (without low value (14 point) bandits from B2023) | | | 0.048* | | 0.067 |
| 0.69 | | 0.57 | | 0.00035* | | 0.0059* | | Equal relative values (all comparisons from all datasets) | | | 0.144 | | 0.159 |

Supplemental Table 3 Quantitative model comparison via the DIC (Spiegelhalter et al. 2014). For tasks p1-p4.2 and the two datasets from Klein et al. 2017 we compared ABS, REL, REP and IER models. For tasks g1, g2 and the datasets from Bavard et al. (2018) and Bavard and Palminteri (2023) we compared ABS, DIV, RAN, RAN$\boldsymbol{\omega}$ and IER models. We further fitted each model twice using two different learning rules. First entry corresponds to context specific learning rates that decay over time (given how often a context was observed) and second entries correspond to model versions with different learning rates for chosen and unchosen options (see methods for details). Further, we fitted an additional version of the REP model, where the repetition bias mechanism was only applied to the learning phase (third entry in the REP column)

| Dataset  (feedback) | DIC(ABS)  learning decay / chosen unchosen | DIC(REL)  learning decay / chosen unchosen | DIC(DIV)  learning decay / chosen unchosen | DIC(RAN)  learning decay / chosen unchosen | DIC(RAN $\boldsymbol{\omega}$)  learning decay / chosen unchosen | DIC(REP)  learning decay / chosen unchosen (REPbias learning only) | DIC(IER)  learning decay / chosen unchosen |
| --- | --- | --- | --- | --- | --- | --- | --- |
| Task p1  (full) | 4562.59/  4584.21 | 4339.53/  4633.33 | - | - | - | 4213.49/  4215.74  (4218.43) | 4562.51/  4580.97 |
| Task p1.1 | 1835.75/  1759.79 | 1732.16/  1811.29 | - | - | - | 1630.17/  1627.91 | 1831.35/  1759.76 |
| Task p1.2 | 1394.21/  1436.62 | 1427.45/  1407.50 | - | - | - | 1389.32/  1329.39 | 1429.45/  1435.32 |
| Task p1.3 | 1278.29/  1319.20 | 1213.24/  1299.91 | - | - | - | 1188.76/  1201.70 | 1278.57/  1318.86 |
| Task p2  (partial) | 2835.23/  2846.81 | 2902.83/  2844.55 | - | - | - | 2696.46/  2633.72  (2697.38) | 2848.28/  2840.97 |
| Task p3  (full) | 3650.67/  3771.47 | 3584.49/  3957.94 | - | - | - | 3503.70/  3493.84  (3457.40) | 3653.67/  3772.71 |
| Task p4.1  (full) | 1794.51/  1739.12 | 1724.28/  1782.94 | - | - | - | 1591.80/  1589.33  (1650.76) | 1796.20/  1736.45 |
| Task p4.2  (full) | 1439.19/  1603.67 | 1364.51/  1590.14 | - | - | - | 1386.69 /  1401.43  (1389.76) | 1438.14/  1603.46 |
| Task g1  (full) | 2595.10/  2305.55 | - | 3602.51/  3030.03 | 3042.93 /  3036.28 | 2264.18/  2386.59 | 2183.07 /  2114.407  (2349.3) | 2074.70/  2047.44 |
| Task g2  (partial) | 4451.66/  4243.97 | - | 5403.30/  5702.21 | 4480.37/  4615.32 | 4243.95/  4506.58 | 4314.60/  4292.12  (4359.31) | 4356.02/  4230.01 |
| Klein et  al. 2017  Exp2  (full) | 1767.75/  1803.455 | 1710.31/  1819.59 | - | - | - | 1527.09/  1506.33  (1534.29) | 1767.71/  1806.93 |
| Klein et  al. 2017  Exp3  (full) | 1542.66/  1549.86 | 1494.54/  1701.26 | - | - | - | 1426.89/  1436.74  (1437.22) | 1555.91/  1536.3 |
| Bavard et  al. 2018  Exp1  (mixed) | 5833.82/  6423.7 | - | 6342.22/  8977.5 | 5913.845/  6634.8 | 5917.86/  6642.5 | 4746.199/  4797.0  (4987.82) | 5390.61/  5990.3 |
| Bavard et  al. 2018  Exp2  (full) | 12146.2/  12384.5 | - | 13234.1/  15055.6 | 11493.73/  11605.1 | 11633.7/  11605.6 | 9601.482/  9754.4  (9537.31) | 10629.8/  11120.7 |
| Bavard et  al. 2023  Exp3a  (full) | 30823.9/  30593.2 |  | 29955.7/  30537.3 | 29039.75/  28041.48 | 28533.3/  28061.1 | 21987.24/  22063.8  (22212.27) | 29062.1/  27785.4 |
| Bavard et  al. 2023  Exp3b  (full) | 31794.3/  32065.4 |  | 32000.9/  33015.5 | 30659.8/  30201.82 | 30205.7/  29432.8 | 24151.11/  24316.81  (25612.4) | 30634.4/  29768.2 |

Supplemental Table 4 Overview of task parametrizations, i.e. reward schedules, number of trials and transfer comparisons for all new and reanalyzed value-based decision tasks. Note for tasks p1-p4.2 and the two data sets from Klein et al. 2017 we only used the first two transfer trials for standard analysis and computational modeling, as these tasks included reward feedback during the tranfer phase.

| New datasets | | | | | | |
| --- | --- | --- | --- | --- | --- | --- |
| Probabilistic tasks | | | | | | |
| Task  (nParticipants) | **Reward probability %**  **(LC context)** | | **reward probability %**  **(HC context)** | **Feedback** | **nTrials**  **learning** | **nTrials**  **transfer (distinct comparisons)** |
| p1.1  (n = 36) | A: 0.7 B: 0.5 | | C: 0.7 D: 0.2 | full | 30 per context | 10 per context  (2 comparisons) |
| p1.2  (n = 30) | A:0.73 B: 0.5 | | C:0.73 D: 0.2 | full | 30 per context | 10 per context  (2 comparisons) |
| p1.3  (n = 29) | A: 0.6 B:0.4 | | C 0.6 D: 0.1 | full | 30 per context | 10 per context  (2 comparisons) |
| p2  (n = 49) | A: 0.7 B: 0.5 | | C: 0.7 D: 0.2 | partial | 30 per context | 10 per context  (2 comparisons) |
| p3  (n = 49) | A: 0.7 B: 0.5 | | C: 0.7 D: 0.4 | full | 50 LC context 30 HC context | 10 per context  (2 comparisons) |
| p4.1  (n = 30) | A: 0.6 B: 0.4 | | C: 0.76 D:0.56 | full | 30 per context | 10 per context  (2 comparisons) |
| p4.2  (n = 29) | A: 0.6 B: 0.4 | | C: 0.76 D: 0.56 | full | 30 per context | 10 per context  (2 comparisons) |
| Gaussian tasks | | | | | | |
| Task  (nParticipants) | **Reward magnitude**  **(LC Context)** | **Reward magnitude**  **(HC Context)** | | **Feedback** | **nTrials**  **learning** | **nTrials**  **transfer** |
| g1  n =49 | A: 70 B: 50 C: 40 | | D: 70 E: 40 F:10 | full | 40 per context | 4 per context (9 comparisons) |
| g2  n = 49 | A: 70 B: 50 C: 40 | | D: 70 E: 40 F:10 | partial | 40 per context | 4 per context (9 comparisons) |

| Existing datasets | | | | | | | | | |
| --- | --- | --- | --- | --- | --- | --- | --- | --- | --- |
| Task | | **Reward probability %**  **LC context** | | | **Reward probability %**  **HC context** | | **Feedback** | **nTrials**  **learning** | **nTrials**  **transfer (distinct comparisons)** |
| Klein et al. 2017 Exp.2  (n = 24) | | A: 0.7 B: 0.5 | | | C:0.7 D: 0.2 | | partial | 30 per context | 30 per context  (1 comparison) |
| Klein et al. 2017 Exp.3  (n = 21) | | A:0.8 B:0.6 | | | C: 0.8 D: 0.6 | | full | 30 per context | 30 per context  (1 comparison) |
| Bavard et al. 2018  Exp.1  (n = 20) | | context 1:  A: 1€ (0.75)  B: 1€ (0.25) | context 2:  C: 0.1€  (0.75)  D: 0.1€ (0.25) | | context 3  E: -1€  (0.25)  F: -1€  (0.75) | context 4  G: -0.1€ (0.25)  H: -0.1€ (0.75) | partial/full | 40 per context | 4 per context  (28 comparisons) |
| Bavard et al.  2018  Exp 2(n = 40) | A: 1(0.75)  B: 1(0.25) | | | C: 0.1(0.75)  D: 0.1(0.25) | E:-1(0.25)  F:-1(0.75) | G:-0.1(0.25)  H:-0.1(0.75) | full | 40 per context | 4 per context  (28 comparisons) |
| Task | | mean  (reward)  Context 1 | mean  (reward)  Context 2 | | mean  (reward)  Context 3 | mean  (reward)  Context 4 |  |  | |
| Bavard and Palminteri 2023  Exp. 3a  (n = 100) | | A : 86  B :50 C :14 | D :86  E :50 F :14 | | G :50  H :32  I :14 | J :50  K :11  L :14 | full (stim D and J blocked in 90 trials) | 45 per context | 2 per context  (66 comparisons) |
| Bavard and Palminteri 2023  Exp. 3b  (n = 100) | | A :86  B :50 C :14 | D :86  E :50 F :14 | | G :50  H :32  I :14 | J :50  K :32  L :14 | full (stim D and J blocked in 135 trials) | 45 per context | 2 per context  (66 comparisons) |

Supplemental Table 5

Supplemental Table 5 Complete pseudorandomized reward sequences for all probabilistic tasks p1.1 to p4.2. Sequences were randomly constructed (except for manual changes at the beginning of task p4.1 and p4.2) with constrains that true reward probabilities should be evident around every 10 to 12 trials and that the last 4 to 5 last trials for the best options in each context are equally rewarded (except for task p1.2 and task 4.2).

| Task | LC/LG Context sequence (reward = 1; no reward = 0) | HC/HG sequence (reward = 1; no reward = 0) |
| --- | --- | --- |
| p1.1 | A_LC_ (0.7):  0 1 1 1 1 1 0 1 0 1 1 0 1 1 1 0 1 1 1 0 0 0 1 1 1 1 1 0 1 1  B_LC_ (0.5):  1 0 0 1 1 1 1 0 1 1 1 0 0 1 1 0 0 0 1 1 0 0 1 1 0 0 0 1 0 0 | C_HC_(0.7):  1 0 1 1 1 0 1 1 1 1 1 1 1 0 1 0 0 0 1 1 0 1 1 0 1 1 1 0 1 1  D_HC_(0.2):  1 0 0 0 0 0 0 1 0 1 0 0 0 0 0 0 0 0 1 1 0 0 0 0 0 0 1 0 0 0 |
| p1.2 | A_LC_(0.73):  0 0 1 1 1 0 1 1 1 1 1 1 1 1 0 1 1 1 1 0 1 1 1 0 1 1 1 1 0 0  B_LC_(0.5):  1 0 1 0 1 0 0 1 1 0 0 0 1 1 1 1 0 0 0 1 0 1 1 0 1 1 0 0 0 1 | C_HC_(0.73):  0 1 1 1 1 1 1 1 1 0 1 1 0 1 1 0 1 1 0 1 1 1 0 0 1 1 1 0 1 1  D_HC_(0.2):  0 0 0 0 0 1 0 0 0 1 1 0 0 0 0 1 0 0 0 0 0 1 0 0 0 0 0 0 1 0 |
| p1.3 | A_LC_(0.6):  1 0 1 0 1 0 1 0 1 1 1 1 0 0 1 0 1 1 1 0 1 1 0 1 1 0 0 1 0 1  B_LC_(0.4):  0 1 0 0 0 1 0 0 1 1 0 1 1 0 0 1 0 1 0 0 1 0 0 1 0 1 0 1 0 0 | C_HC_(0.6):  0 1 1 0 0 1 1 1 0 1 0 1 0 1 1 0 1 1 1 0 1 0 1 0 1 1 0 1 0 1  D_HC_(0.1):  0 0 0 0 1 0 0 0 0 0 0 0 0 0 1 0 0 0 0 0 0 0 1 0 0 0 0 0 0 0 |
| p2 | A_LC_(0.7):  0 1 1 1 1 1 0 1 0 1 1 0 1 1 1 0 1 1 1 0 0 0 1 1 1 1 1 0 1 1  B_LC_(0.5):  1 0 0 1 1 1 1 0 1 1 1 0 0 1 1 0 0 0 1 1 0 0 1 1 0 0 0 1 0 0 | C_HC_(0.7):  1 0 1 1 1 0 1 1 1 1 1 1 1 0 1 0 0 0 1 1 0 1 1 0 1 1 1 0 1 1  D_HC_(0.2):  1 0 0 0 0 0 0 1 0 1 0 0 0 0 0 0 0 0 1 1 0 0 0 0 0 0 1 0 0 0 |
| p3 | A_LC_(0.7):  1 1 0 1 0 1 1 1 0 1 0 1 1 1 0 1 1 1 1 0 0 0 1 0 1 1 1 1 1 1 1 0 1 1 0 1 0 1 1 1 1 1 0 0 1 0 1 1 1 1  B_LC_(0.5):  1 0 0 1 0 1 1 1 1 0 1 1 1 0 0 1 1 0 0 0 0 0 0 1 0 1 1 1 0 0 1 1 0 1 1 1 0 0 0 1 0 1 0 1 1 0 0 1 0 0 | C_HC_(0.7):  0 1 1 0 1 1 1 0 1 1 1 0 1 0 1 1 1 1 0 1 0 1 1 0 1 0 1 1 1 1  D_HC_(0.4):  1 0 0 0 1 0 1 0 0 1 0 1 0 1 1 0 0 1 0 0 0 1 0 1 1 0 0 1 0 0 |
| p4.1 | A_LG_(0.6):  1 1 0 1 0 1 0 1 1 0 0 1 1 0 1 0 1 0 1 0 1 0 1 1 1 0 1 0 1 1  B_LG_(0.4):  1 1 0 0 1 0 0 0 1 0 0 0 1 0 1 0 0 1 0 1 0 1 0 1 0 1 0 1 0 0 | C_HG_(0.76):  0 1 1 1 1 1 0 1 1 1 1 0 1 1 1 0 1 1 1 0 1 0 1 1 1 1 1 0 1 1  D_HG_(0.56):  1 0 0 1 1 1 1 0 1 1 1 0 0 1 1 0 0 1 1 1 0 0 1 1 0 1 0 1 0 0 |
| p4.2 | A_LG_(0.6):  0 1 0 1 0 1 0 1 1 0 0 1 1 0 1 0 1 1 1 0 1 0 1 1 1 0 1 0 1 1  B_LG_(0.4):  1 1 0 0 1 0 0 0 1 0 0 0 1 0 1 1 0 1 0 0 1 1 0 0 1 0 0 1 0 0 | C_HG_(0.76):  1 1 1 1 1 1 0 1 1 0 1 0 1 1 1 0 0 0 1 1 1 1 1 1 1 1 1 0 1 1  D_HG_(0.56):  0 0 0 1 1 1 1 0 1 1 1 0 0 1 1 0 1 0 1 1 1 1 0 1 0 0 1 1 0 0 |

*Repetition bias*

In the REP model, the repetition learning rate parameter $\eta_{\mathrm{repetition}}$ primarily ranged from 0.03 to 0.25 across 12 of the 15 analyzed tasks (see Supplemental Table 7 for summary statistics and Supplemental Figure 1 for visualization). Although this parameter varied with different task parameterizations, it remained relatively consistent across participant groups performing identical tasks under the same conditions (see Supplemental Figures 1B, 1D, and 1E). We interpret this stability as evidence of a common underlying mechanism driving choice behavior across diverse task dynamics.

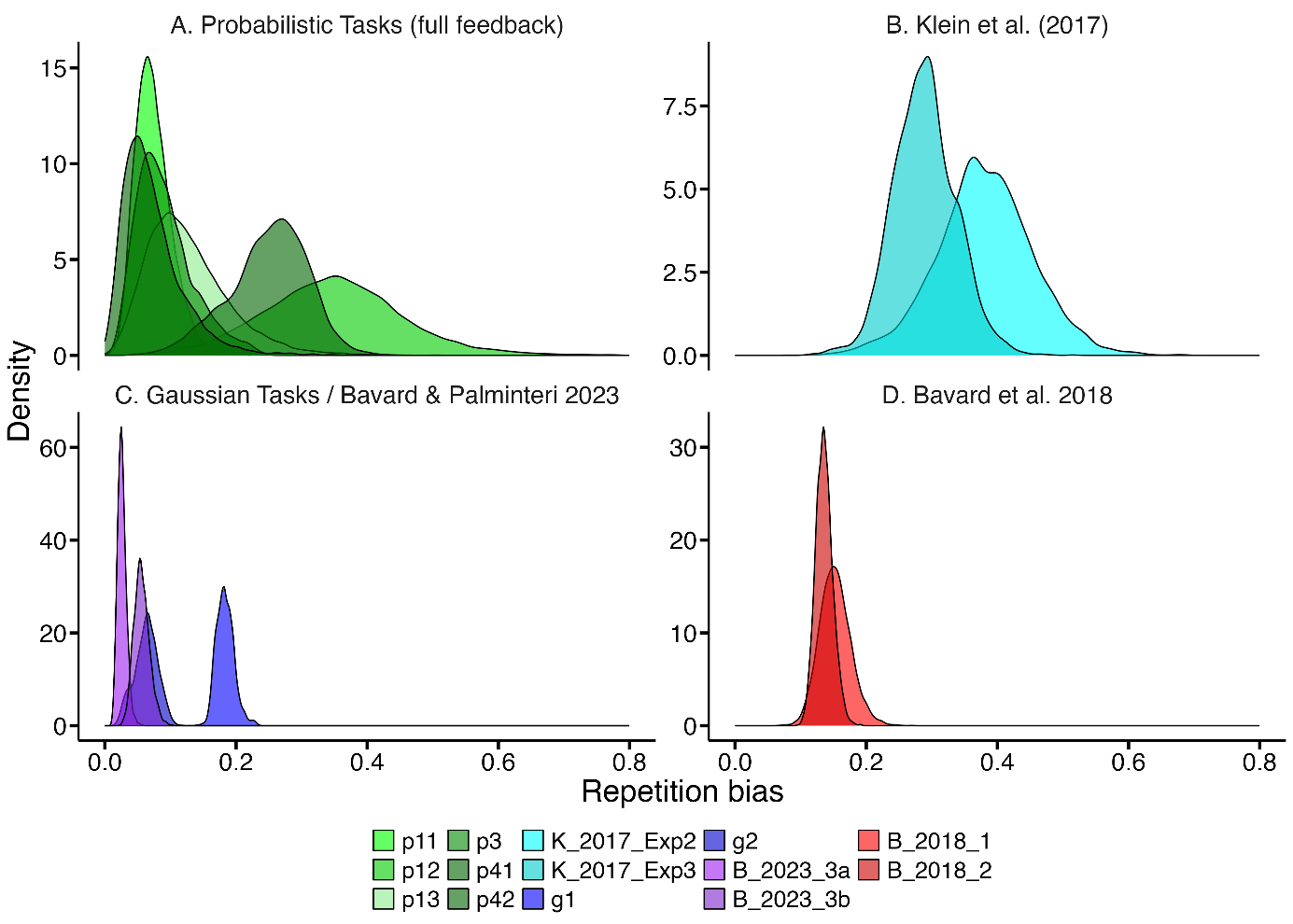

Supplemental Figure 1 $\eta_{repetition}$ varies between different datasets but remains relatively stable among participant groups performing identical tasks. For example, most parameter estimates for probabilistic tasks range from 0.05 to 0.2. Tasks from Klein et al. (2017) exhibit substantial overlap in their $\eta_{repetition}$ hyperparameter distributions, while datasets from Bavard and Palminteri (2023) and Bavard et al. (2018) display similar repetition-bias parameters (see Supplemental Table 7 and Supplemental Figure 1 for details).

Supplemental Table 6 95% highest density interval (HDI) and mean values for $\eta_{repetition}$ hyperparameter distributions across all analyzed datasets.

| Task | p1 | p2 | p3 | p4.1 | p4.2 | Klein et al. 2017  Exp2 | Klein et al. 2017 Exp3 |
| --- | --- | --- | --- | --- | --- | --- | --- |
| 95% HDI (lower-upper) | 0.0959-  0.2535 | 0.0013-0.0196 | 0.0207-  0.1809 | 0.1270-  0.3579 | 0.0029-  0.1595 | 0.2459-  0.5389 | 0.1997-  0.3834 |
| Mean | 0.1722 | 0.0064 | 0.0915 | 0.2498 | 0.0726 | 0.3810 | 0.2891 |
| Task | **g1** | **g2** | **Bavard et al. 2018 Exp1** | **Bavard et al. 2018 Exp2** | **Bavard and Palminteri 2023 Exp3a** | **Bavard and Palminteri 2023 Exp 3b** |  |
| 95% HDI (lower-upper) | 0.1582-  0.2098 | 0.0264-  0.0947 | 0.1067-  0.2003 | 0.1105-  0.1614 | 0.0143-  0.0391 | 0.0347-  0.0804 |  |
| Mean | 0.1835 | 0.0630 | 0.1525 | 0.13597 | 0.02601 | 0.0561 |  |

*Different learning mechanisms*

We compared two distinct learning mechanisms in our study. First, following Sutton and Barto (2018), we implemented decaying learning rates, which are appropriate when reward probabilities (bandits) are stationary. Second, consistent with prior research, we fitted separate constant learning rates for chosen and unchosen options. Overall, models with context-specific decaying learning rates outperformed models with different learning rates for chosen and unchosen options in 30 out of 40 models applied to probabilistic task data and in 9 out of 24 models applied to Gaussian task data (see Supplemental Table 3 for an overview).
